# Automated Extraction and Visualization of Metabolic Networks from Biomedical Literature Using a Large Language Model

**DOI:** 10.1101/2023.06.27.546560

**Authors:** Thiptanawat Phongwattana, Jonathan H Chan

## Abstract

The rapid growth of biomedical literature presents a significant challenge for researchers to extract and analyze relevant information efficiently. In this study, we explore the application of GPT, the large language model to automate the extraction and visualization of metabolic networks from a corpus of PubMed abstracts. Our objective is to provide a valuable tool for biomedical researchers to explore and understand the intricate metabolic interactions discussed in scientific literature. We begin by splitting a ton of the tokens within the corpus, as the GPT-3.5-Turbo model has a token limit of 4,000 per analysis. Through iterative prompt optimization, we successfully extract a comprehensive list of metabolites, enzymes, and proteins from the abstracts. To validate the accuracy and completeness of the extracted entities, our biomedical data domain experts compare them with the provided abstracts and ensure a fully matched result. Using the extracted entities, we generate a directed graph that represents the metabolic network including 3 types of metabolic events that consist of metabolic consumption, metabolic reaction, and metabolic production. The graph visualization, achieved through Python and NetworkX, offers a clear representation of metabolic pathways, highlighting the relationships between metabolites, enzymes, and proteins. Our approach integrates language models and network analysis, demonstrating the power of combining automated information extraction with sophisticated visualization techniques. The research contributions are twofold. Firstly, we showcase the ability of GPT-3.5-Turbo to automatically extract metabolic entities, streamlining the process of cataloging important components in metabolic research. Secondly, we present the generation and visualization of a directed graph that provides a comprehensive overview of metabolic interactions. This graph serves as a valuable tool for further analysis, comparison with existing pathways, and updating or refining metabolic networks. Our findings underscore the potential of large language models and network analysis techniques in extracting and visualizing metabolic information from scientific literature. This approach enables researchers to gain insights into complex biological systems, advancing our understanding of metabolic pathways and their components.

## Introduction

The field of biomedical research is experiencing an unprecedented surge in the volume of scientific literature, presenting researchers with a formidable challenge of extracting and analyzing relevant information efficiently. In this groundbreaking study, we delve into the transformative application of the GPT-3.5-Turbo language model to automate the extraction and visualization of metabolic networks from a carefully curated corpus of 18 PubMed abstracts from relevant prior work. The primary objective of our research is to provide a powerful and indispensable tool for biomedical researchers, facilitating their exploration and comprehension of the intricate metabolic interactions extensively discussed within the vast realm of scientific literature.

To embark on this ambitious endeavor, we initiate the process by meticulously splitting the testing corpus into two distinct corpora. This strategic division is necessitated by the inherent limitations of the GPT-3.5-Turbo model, which imposes a token limit of 4,000 per analysis. With unwavering dedication and meticulous attention to detail, we embark on an iterative prompt optimization process. This iterative refinement process plays a pivotal role in the successful extraction of a comprehensive list of metabolites, enzymes, and proteins from the abstracts, ensuring the highest degree of accuracy and completeness in the obtained entities.

With the extracted entities at our disposal, we embark on the awe-inspiring task of generating a visually captivating and highly informative directed graph. This monumental achievement is accomplished through the strategic fusion of Python programming and the versatile NetworkX library. The resulting graph visualization serves as an awe-inspiring testament to the immense power that emerges from the seamless integration of cutting-edge language models and sophisticated network analysis techniques. It offers a lucid and comprehensive representation of the intricate metabolic pathways that underscore the delicate balance and interplay between metabolites, enzymes, and proteins. This transformative approach showcases the immense potential of automated information extraction and advanced visualization methodologies in shedding light on the enigmatic world of metabolic interactions.

The contributions of our research endeavor are not limited to a singular facet; rather, they are twofold, each bearing profound significance. Firstly, we take great pride in presenting a groundbreaking demonstration of the exceptional prowess of the GPT-3.5-Turbo language model, as it deftly and autonomously extracts a vast array of metabolic entities. By streamlining the arduous process of cataloging and discerning important components in the realm of metabolic research, our research showcases the immense potential of large language models in catapulting scientific inquiry to new heights. By harnessing the vast capabilities of the GPT-3.5-Turbo language model, we empower researchers to transcend the constraints of manual extraction, thus paving the way for a more efficient and accelerated pace of scientific progress in the field of metabolic research.

Secondly, we present a momentous innovation in the generation and visualization of a meticulously crafted directed graph. This visually captivating graph provides a comprehensive and all-encompassing overview of the intricate metabolic interactions that underpin the vast array of biological systems. It serves as an invaluable tool, offering researchers a robust framework for further analysis, meticulous comparison with existing pathways, and iterative refinement of metabolic networks. The transformative potential of this graph lies in its ability to unlock hidden patterns, unravel the enigmatic web of connections between metabolites, enzymes, and proteins, and provide researchers with a solid foundation for advancing our understanding of the complex world of metabolism.

The implications of our research findings reverberate throughout the scientific community, resonating with the immense potential that lies dormant within large language models and network analysis techniques. Our research endeavors serve as a clarion call, heralding the transformative potential of these innovative methodologies in the extraction and visualization of metabolic information gleaned from the vast expanse of scientific literature. By bridging the gap between automated information extraction and sophisticated visualization methodologies, we equip researchers with an invaluable toolkit that propels scientific inquiry to unprecedented heights. Our research not only fuels breakthrough discoveries but also catalyzes transformative advancements in the field of biomedical informatics, thereby accelerating our understanding of the intricate metabolic pathways and their constituent components. Through this pioneering research, we usher in a new era of knowledge exploration and scientific progress.

### Related Work

Preecha P., et al. [1] presents an integrated TM framework consisting of two modules: the Metabolic Event Extraction module (MEE) and the Metabolic Interaction Network Reconstruction module (MINR).

The integrated TM framework demonstrates promising performance, as evaluated based on recall, precision, and F-score measures. The MEE module achieves F-scores of 59.15% and 48.59% for detecting metabolic events related to production and consumption, respectively. The entity tagger for Gene and Protein (GP) and metabolite achieves an F-score greater than 80% for the Superpathway of leucine, valine, and isoleucine biosynthesis. The MINR module, which maps enzyme and metabolite interactions, achieves an F-score exceeding 70% on the test corpus.

Furthermore, the integrated TM framework is applied to a large-scale dataset (EcoCyc extraction data) for reconstructing a metabolic interaction network. The reconstruction shows reasonable precision for enzyme (69.93%), metabolite (70.63%), and enzyme-metabolite (46.71%) interactions. Notably, this study presents the first open-source integrated TM framework for reconstructing metabolic interaction networks, providing biologists with a powerful tool for extracting metabolic events and reconstructing such networks.

An event extraction system that utilizes text mining techniques to extract metabolic interactions from research literature and reconstruct metabolic pathways. The proposed system from previous study [2] consists of a four-step pipeline: named entity recognition, trigger detection, edge detection, and event reconstruction. To enhance the classification of metabolic event extraction in the target domain, the authors introduce a multitask-learning algorithm that leverages additional resources from an existing source domain, following a transfer-learning paradigm.

As a proof of concept, the authors focus on edge detection as a core step in the event extraction system and conduct a case study using multitask-learning classification. Experimental results demonstrate that the proposed event extraction system achieves competitive performance compared to state-of-the-art systems. Notably, the multitask-learning approach improves the performance of edge detection, subsequently enhancing the overall performance of the event extraction system.

This work contributes to the advancement of text mining techniques for extracting metabolic pathways from research literature. By introducing an event extraction system with a multitask-learning algorithm, the authors demonstrate the potential for automated extraction and reconstruction of metabolic interactions. This research highlights the importance of text mining in facilitating the analysis of biological data and provides a valuable tool for biologists in their quest for extracting relevant information from unstructured text.

BioBERT is a pre-trained language representation model specifically designed for the biomedical domain. It is trained on a large corpus of biomedical text data, including scientific articles, medical records, and other relevant documents, to capture the knowledge and language patterns specific to the biomedical field. The goal of BioBERT is to provide a highquality representation of biomedical text data that can be fine-tuned for various NLP tasks, such as named entity recognition, sentiment analysis, and question answering [3]. There is also a biomedical data study addresses the challenge of accurate unknown entity recognition in the context of NER. By employing the BERT model for NER classification and leveraging additional biomedical corpora, the study achieved significant improvements in protein recognition within the specific super-pathway of interest. Furthermore, the development of an internal web platform demonstrates practical applicability, supporting bio-annotators and enabling further corpus enlargement efforts [4].

PubMed, on the other hand, is a publicly accessible database of biomedical literature maintained by the National Library of Medicine (NLM) at the National Institutes of Health (NIH) [5], [6]. It contains over 30 million citations and abstracts of scientific articles, medical papers, and other related content in the biomedical domain, including genetics, pharmacology, and more. PubMed is widely used by researchers, healthcare professionals, and the general public as a resource for finding and accessing scientific literature in the biomedical field. BERT (Bidirectional Encoder Representations from Transformers) uses a training technique called “Masked Language Modeling” to train its model. The idea behind masked language modeling is to randomly mask some of the words in a sentence, and then train the model to predict the original words based on the context [7].

In more detail, during training, the BERT model [8] is given a sentence, and a percentage of the words in the sentence are masked (e.g., 15% of the words are masked). The masked words are represented as special tokens (e.g., [MASK]). The model then uses the context of the unmasked words to predict the original words of the masked tokens. This task requires the model to understand the context of the sentence and the relationships between words in the sentence, which in turn helps it learn a contextualized representation of the words in the sentence [4].

BERT algorithm utilizes attention mechanism with the following formula.

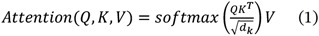

Q is a matrix that represents a dimensional vector of each word in the sequence.

K is a key that represents dimensional vectors of all the words in the sequence.

V is a value that represents dimensional vectors, which are matched to K’s keys.

As in the abovementioned, the formula can be simplified (1) as below.

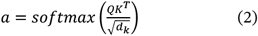

By masking words randomly and training the model to predict the original words, BERT can learn a general representation of the language that can be fine-tuned for various NLP tasks, such as named entity recognition, sentiment analysis, and question answering. The ability of BERT to learn contextual relationships between words in a sentence is what sets it apart from traditional language representation models and makes it a powerful tool for NLP. was used for our training and the word embedding architecture is described as in figure 2.

**Figure 1:**
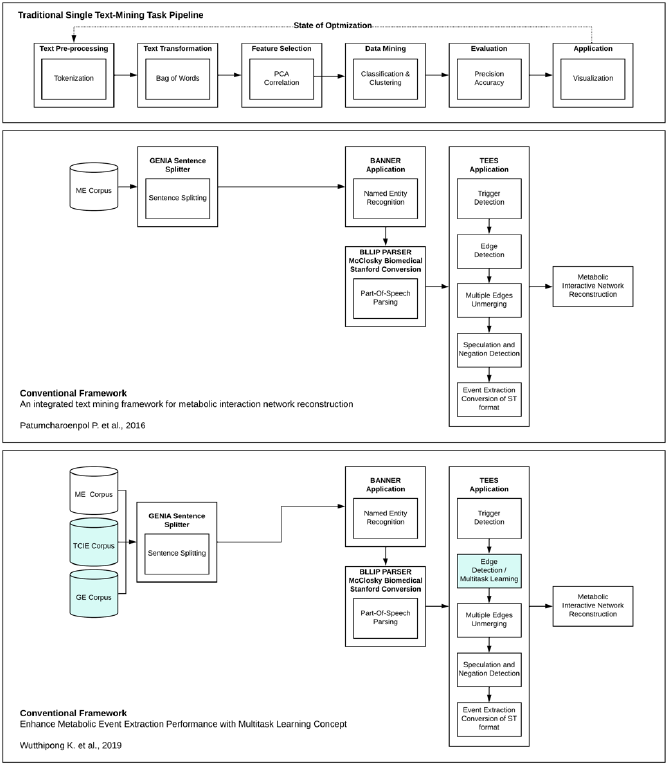
The related work of metabolic interaction network reconstruction by using a text mining technique [1], [2].

**Figure 2:**
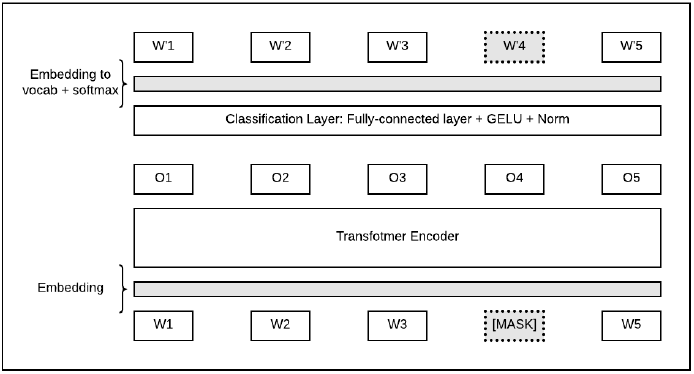
Bidirectional Encoder Representations from Transformers in a masked-word prediction.

As shown in Figure 2, which represents a high-level architecture of the BERT masking process for predicting a sequence of tokens. The size of input and output is equivalent. The loss function comprises of only the computation of the masked values prediction and ignores the non-masked words..

In the study [8], as shown in Figure 3, the authors use a pair of sentences as an input. All sentences will be divided into two sets, each being 50%. The first half includes pairs of one sentence and its subsequent sentence within the original document; the second half includes pairs of a sentence and a random sentence that is disconnected from the first sentence, within the original document.

**Figure 3:**
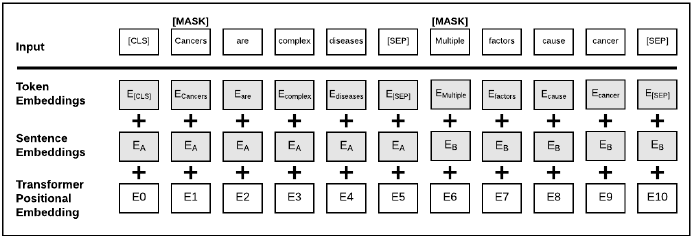
Modified BERT with multiple biomedical sentences representation.

**Figure 4:**
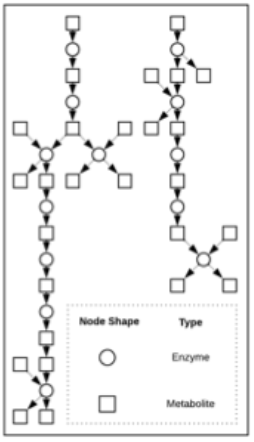
Super-pathway of leucine, valine, and isoleucine biosynthesis [1].

To recognize the first sentence in each input, a [CLS] token is inserted at the first position of the sentence. To recognize the end of each sentence, a [SEP] token inserted at the end. A sentence embedding is also added to each token. That means all tokens in one sentence will have the same sentence embedding.

A positional embedding is also needed within the architecture for defining the position of each token within the sequence. This technique is the main part of the Transformer model.

The pre-training of models on large scale unlabeled data and then fine-tuning them for downstream tasks has proven to be an effective strategy in deep learning. The pre-trained models provide a foundation of learned representations that can be utilized to enhance the performance of the down-stream tasks.

In the field of biomedicine, it has been established that fine-tuning pre-trained language models on in-domain data can significantly improve performance on downstream tasks. Various studies have attempted to improve the performance of BERT models in the biomedical domain by pre-training on specific biomedical text data, such as PubMed abstracts and full-text articles, as well as clinical notes from the MIMIC-III database [9]. However, BERT, which learns powerful word representations through a bi-directional Transformer encoder model architecture, has limitations in its ability to generate text.

To address this limitation, Generative Pre-trained Transformer (GPT) [10] has become promising. GPT was introduced as a solution for language generation tasks. The model was pre-trained on a large corpus of text data using a classical casual language modeling task, where it learns to predict the next word token based solely on the previous word tokens. Furthermore, the development of GPT-2 and GPT-3, with larger model sizes pre-trained on even larger text corpora, resulted in remarkable performance in various down-stream tasks such as translation, summarization, and even classification tasks like reading comprehension, without the need for fine-tuning, through the careful design of appropriate prompts. However, original GPT models have shown poor performance on biomedical tasks despite their powerful generation capabilities. Previous works, such as DARE, which pre-trained GPT on a limited amount of data (0.5 million PubMed abstracts) for use in data augmentation for relation extraction, and a recent work using GPT for unconventional clinical tasks, have not produced satisfactory results. Whereas pre-training BERT model from scratch on large corpora of scientific literature, mainly in the biomedical and computer science domains, is a more effective strategy. This approach allows for a more suitable vocabulary that is better suited to the biomedical domain, leading to the creation of models such as PubMedBERT [11], which is pre-trained on 14 million PubMed abstracts, and the more advanced ELECTRA model, pre-trained on 28 million data.

For the downstream tasks, they can be categorized into sub-sections as follows.

- Relation extraction is a crucial task for the biomedicine and life science fields. Conventional pipeline-based approaches treat the task as a sequence of separate sub-tasks that require intermediate annotations and information, which may suffer from a lack of annotated data and error accumulation.
- Joint extraction, on the other hand, aims to extract both entities and the relationships between them from the text in a coordinated manner. Sequence labeling methods address the task by labeling each word token in the text with different tags to identify entity mentions, and then classify the relationships between them using classifiers.
- Table filling methods, in contrast, formulate the task as a table created through the Cartesian product of itself, and predict the relationships between token pairs. These methods may also suffer from error accumulation due to the previous tagging process and the need for extensive intermediate annotations, such as named entity recognition.
- Text generation methods instead view the task as a sequence-to-sequence learning task, where the input is the text and the target is a triplet, and an encoder-decoder network is employed to learn how to generate the triplet from the text. However, many joint extraction methods still require supplementary entity information.
- Question answering (QA) is a task in which the goal is to provide an answer to a question based on a given context. This task has been approached in various ways, including predicting a span within the source context as the answer text or predicting a label (e.g., “yes” or “no”) for simpler tasks with predefined categorical answers. The field of biomedical domain question answering has seen a significant amount of research utilizing pre-trained language models. Generative models have been employed to directly generate the answer sequence or label words.

In comparison, traditional models for biomedical event extraction often rely on hand-crafted features, such as part-of-speech tags, named entity recognition, and dependency parsing. These models are trained from scratch on labeled data and do not have the pre-training and transfer learning capabilities of GPT-3. While these traditional models can still achieve good results on biomedical event extraction tasks, they typically require much larger amounts of labeled data and may not be as effective at capturing the complex relationships and context in biomedical text.

A recent study has proposed BioGPT [12], a domain-specific Generative Pre-trained Transformer (GPT) language model designed for biomedical text generation and mining. The model is based on the Transformer architecture and was pretrained on 15 million PubMed abstracts from scratch. The effectiveness of BioGPT was evaluated on six different biomedical natural language processing (NLP) tasks, including end-to-end relation extraction on BC5CDR, KD-DTI, and DDI datasets, question answering on the PubMedQA dataset [13], document classification on the HoC dataset, and text generation.

There is a study that analyzed the target sequence format and the prompt for each task to optimize the performance of BioGPT. The results of the experiments showed that BioGPT outperformed the baseline models on all the NLP tasks. The evaluation of clinical knowledge in models is currently limited to automated evaluations on limited benchmarks. There is a lack of standard for evaluating model predictions and reasoning across a range of tasks. In order to address this issue, a study was conducted that proposed MultiMedQA [14], a benchmark that integrates six existing open question answering datasets covering various medical exams, research and consumer queries, and HealthSearchQA [14], a new free-response dataset of medical questions searched online.

The study also presented a framework for human evaluation of model answers along several dimensions including factuality, precision, potential harm and bias. The study evaluated PaLM [15](a large language model with 540 billion parameters) and its instruction-tuned variant, Flan-PaLM, on MultiMedQA. Through the use of various prompting strategies, Flan-PaLM achieved state-of-the-art accuracy on all the multiple-choice datasets in MultiMedQA (MedQA, MedMCQA, PubMedQA, and MMLU clinical topics), including 67.6% accuracy on MedQA (US Medical License Exam questions), outperforming the previous state-of-the-art by more than 17%. However, human evaluation revealed some limitations in the answers provided by Flan-PaLM. To overcome this, the study introduced instruction prompt tuning, an efficient approach for aligning large language models to new domains using a small number of examples. The resulting model, Med-PaLM, showed promising results, but still fell short of clinician performance. The study highlights that model scale and instruction prompt tuning can improve comprehension, recall of knowledge, and medical reasoning. The human evaluations also emphasize the importance of both evaluation frameworks and method development in creating safe and helpful large language models for clinical applications.

In the field of medical research, systematic reviews are widely regarded as the most comprehensive form of literature review to address a focused research question. These reviews are often considered the gold standard in evidence-based medicine and play a critical role in answering questions in the medical field. The creation of a high-quality systematic review requires the construction of complex Boolean queries to retrieve relevant studies for the review topic. However, the process of constructing these queries is often time-consuming and may result in an ineffective query, which can lead to biased or invalid reviews. This can occur when key evidence is missed or when too many irrelevant studies are retrieved, leading to increased review costs.

Advances in Transformer-based generative models have shown great promise in effectively following user instructions and generating answers based on those instructions. A recent study investigated the effectiveness of one such model, ChatGPT [16], in generating effective Boolean queries for systematic review literature searches. Through a series of comprehensive experiments on standard test collections for this task, the study found that ChatGPT could generate queries that result in high search precision, even if recall was traded off. The study highlights the potential of ChatGPT in generating effective Boolean queries for systematic review literature searches and its ability to follow complex instructions and generate queries with high precision makes it a valuable tool for researchers conducting systematic reviews, particularly for rapid reviews where time is a constraint and a trade-off between precision and recall may be acceptable.

According to a recent study, the authors explored the use of graph neural network (GNN) techniques [17] for predicting links within biomedical networks [18]. Biomedical networks, which model the interactions between various entities such as proteins and drugs, play a crucial role in understanding the pathological mechanisms of human diseases and in drug discovery. Link prediction in these networks is a challenging task, as it requires the integration of multiple data sources, including sequence, structure, and network data, to effectively extract features for each node.

To address this challenge, the authors proposed a novel pretraining model that integrates different data sources for link prediction in biomedical networks. The model leverages expressive deep learning methods, such as Convolutional Neural Networks (CNNs) [19], [20] and Graph Convolutional Networks (GCNs) [18], [21], to extract features from sequence and structure data. The GCN-based encoder then refines these features by modeling the dependencies among nodes in the network data. The pre-training model is based on graph reconstruction tasks, and its effectiveness was evaluated on two critical link prediction tasks: synthetic lethality prediction and drug-target interaction prediction.

As in the experimental results, it showed that the features generated by the pre-training model could significantly improve the performance and reduce the training time for existing GNN models. Furthermore, fine-tuning the pretrained model to a specific task could achieve performance comparable to state-of-the-art methods. In addition, the study also highlights the potential of pre-training models in integrating multiple data sources for link prediction in biomedical networks.

In recent years, the outbreak of Coronavirus Disease-19 (COVID-19) [22] has caused a global epidemic with significant morbidity and mortality. Despite extensive research efforts, there is currently no proven effective treatment for COVID-19. The identification of drug-virus associations is crucial in understanding the underlying mechanisms of drug-virus interaction and in guiding the screening process for antiviral drug candidates. Conventional experimental methods for this purpose are often time-consuming, labor-intensive, and costly. Thus, computational approaches provide a valuable alternative for identifying potential drug candidates for viruses, such as COVID-19. In this regard, a study has proposed a framework for drug-virus association prediction using Heterogeneous Graph Attention Networks, named HGATDVA [23]. This framework provides a novel approach for identifying potential drug candidates for COVID-19 and other viruses through computational means.

The representation of metabolic pathways as graphs provides the opportunity for advanced analysis. The utilization of Neo4j [24], a graph database management system, has been demonstrated in several studies for the visualization of metabolic pathway data, commonly referred to as ‘Recon2Neo4j’ in the literature. Recon2Neo4j is a representation of the metabolic pathways data in the form of a graph, using the graph database technology Neo4j. It provides a way to visualize and analyze metabolic pathways data using Neo4j. For example, Recon2Neo4j can be used to analyze the relationships between different metabolites and enzymes in a metabolic pathway, as well as the flow of metabolites through the pathway. This information can be useful for understanding the underlying biological mechanisms involved in metabolic pathways, and for identifying potential targets for drug discovery. Another example of the use of Recon2Neo4j is in the analysis of large-scale metabolic pathway data, such as data from omics studies. By visualizing the data in a graph format, it becomes easier to identify patterns and relationships in the data, and to gain insights into the metabolic pathways that are most relevant to a given biological system.

Several research efforts have been made in the field of finding biosynthetic pathways and analyzing metabolic networks. In this section, we discuss relevant works that address the challenges of pathway extraction, network analysis, and efficient graph search algorithms. One prominent challenge in metabolic engineering and bioproduction pathway elucidation is the extraction of novel metabolic pathways from large-scale networks. With the integration of big data, the efficient analysis and navigation of metabolic networks have become increasingly important. The proposed approach by [Author1] addresses this challenge by constructing searchable graph representations of metabolic networks. They decompose each reaction into pairs of reactants and products and assign weights based on the number of conserved atoms between them. This method has shown promising results in extracting biologically relevant metabolic pathways from biochemical networks [25].

## Methodology

In this study, we use GPT-3.5 Turbo model that is based on Instruction-Tuned LLM (Language Model) to extract bio-entities and relationships from each abstract that is fetched from PubMed via Entrez API. Instruction-Tuned LLM refers to a methodology that involves incorporating carefully designed instructions during the fine-tuning stage of a language model. LLMs are pre-trained models that learn the statistical patterns and structures of language through exposure to a vast corpus of text data. However, their behavior and performance can be further optimized for specific tasks or domains by providing explicit instructions during the fine-tuning process. The instructions in Instruction-Tuned LLM serve as guiding signals that direct the model’s behavior and shape its responses to specific inputs. These instructions can take various forms, such as prompts, demonstrations, or constraints, and are designed to align the model’s output with desired outcomes. By explicitly instructing the model on how to respond or reason, researchers can enhance the accuracy, relevance, and contextuality of the model’s outputs. The design of effective instructions requires a deep understanding of the target task or domain, as well as the capabilities and limitations of the language model. Instructions should be clear, specific, and relevant to the desired outcome, providing guidance without overly constraining the model’s flexibility.

Instruction-Tuned LLM offers several advantages. First, it allows for task-specific optimization, enabling researchers to fine-tune the model’s behavior to meet specific requirements. We utilize these advantages for finding bio-entities and relationships. We found that we must define task-specific prompts for well-recognizing and yielding best results. This can lead to improved performance and more accurate results in the target task.

Second, Instruction-Tuned LLM supports domain adaptability by incorporating domain-specific instructions during fine-tuning. According to this adaptability, we provide a prefix for all prompts that define the model as a biomedical data expert. This helps the model generate more relevant and accurate responses in specialized domains or narrow contexts. Third, instructions can be used to address ethical concerns and mitigate biases in language models. By explicitly instructing the model to prioritize fairness, inclusivity, or avoidance of sensitive topics, Instruction-Tuned LLM contributes to reducing biased outputs and promoting responsible AI use.

We leverage LLM instructions to guide the behavior of language models. By doing so, Instruction-Tuned LLM offers a valuable approach to enhancing their capabilities and aligning them with specific tasks we assign for metabolic network domain.

### Prompt Optimization Methodology

For improving responses from complex tasks such as Biomedical domain, iterative prompt development is required. The process that involves refining and improving prompts or instructions provided to a language model has the exact goal which is to iteratively enhance the model’s responses and performance by fine-tuning the prompts through an iterative feedback loop. And it can be described as below.

#### 1. Initial Prompt Design

In the first iteration, an initial set of prompts or instructions is designed based on the desired task or objective. These prompts should be clear, specific, and aligned with the intended outcome.

You are a biomedical data expert. \

There are 2 main tasks for you and you need to make sure that a final result you \ provide is completely based on my information, not from other sources. \

Task 1: the task is to recognize biomedical entities from the provided text that consist of metabolites, enzymes, \and proteins into a table with 2 headers that consist of “Entity Name” and “Entity Type”. \

Task 2: the task is to recognize metabolic event types from the provided text that consist of metabolic_reaction, \ metabolic_consumption, metabolic_production. And display its relevant entities in a table \ with 3 headers that consist of “Event Name”, “Event Type”, and “Relevant Entity” from Task 1 for each event.

#### 2. Model Fine-tuning

The initial prompts are used to fine-tune the GPT model on a specific task or dataset. This involves training the model on the task-specific data while considering the provided prompts as guiding signals for generating appropriate responses. In this study we utilize a process of natural language processing (NLP) that is based on large language models to PubMed abstracts. By comparing to the prior work’s NLP pipeline, it is not necessary to follow the step-by-step instruction as in Figure 5. As a result, LLM properly works as the end-to-end process in natural language processing.

**Figure 5:**
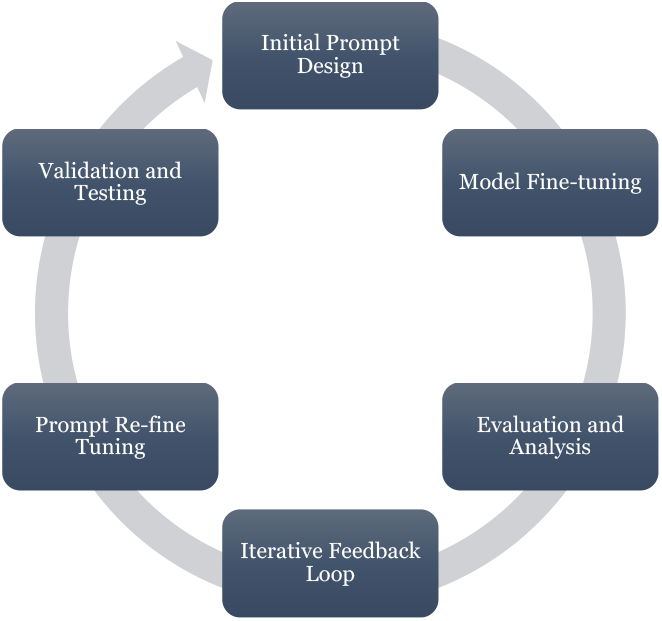
Prompt Optimization Cycle.

#### 3. Evaluation and Analysis

The fine-tuned model’s outputs are evaluated to assess its performance. This evaluation can involve manual review, automated metrics, or a combination of both. The objective is to analyze the model’s strengths, weaknesses, and areas for improvement. In our study, we have domain experts to evaluate manually. We evaluate by using EcoCyc metabolic pathway as the ground truth, and also compare with the current baseline from the relevant previous work.

#### 4. Iterative Feedback Loop

Based on the evaluation and analysis, the prompts are refined and modified to address the identified issues or enhance specific aspects of the model’s performance. This can involve adjusting the wording, changing the format, or introducing additional instructions to guide the model’s behavior. In order to iteratively optimize existing prompts, we need to change some words in the study and deliberately validate responses to ensure that the responses are not hallucinated that are abstractedly generated by LLMs.

#### 5. Prompt Re-Fine tuning

The refined prompts are used to fine-tune the model again, incorporating the updated instructions. This iterative process is repeated several times to iteratively improve the model’s responses and optimize its behavior for the target task.

#### 6. Validation and Testing

The final fine-tuned model with the iteratively developed prompts is validated and tested on a separate evaluation corpus to ensure its improved performance and generalization for any further abstracts from PubMed that the system automatically collects via Entrez API.

The key advantage of iterative prompt development is the ability to fine-tune the model based on continuous feedback and analysis. By refining and adapting the prompts over multiple iterations, we found that it can gradually align the model’s responses with the desired outcome, improving its accuracy, contextuality, and relevance.

This approach allows for a more nuanced and tailored optimization of the language model, as we can address specific issues, biases, or limitations that may arise during the initial fine-tuning process in our metabolic pathway information extraction in both its entities and relationships. It also provides an opportunity to explore the interplay between prompts and model behavior, allowing for the discovery of effective instruction strategies.

Our study was conducted using a combination of literature analysis and computational modeling. We began by gathering information from various sources, including scientific publications and databases, to identify key metabolites, proteins, enzymes, and other relevant components involved in metabolic processes. This information served as the foundation for constructing a comprehensive table that outlined the nodes and their corresponding types.

With the table in place, we utilized the NetworkX library in Python to create directed graphs representing the metabolic interactions. The nodes were assigned types based on their categorization, such as metabolite, enzyme, or protein. The edges, representing metabolic interactions, were classified into three types: metabolic_consumption, metabolic_reaction, and metabolic_production. By mapping the relationships between the nodes and the edges, we constructed a visual representation of the metabolic network. For the methodology, it can be described as follows.

### Data Collection

The first step in our methodology was gathering data from diverse sources, including scientific publications and databases. This data collection process aimed to identify key metabolites, proteins, enzymes, and other relevant components involved in metabolic processes. We carefully curated this information to construct a comprehensive table that served as the foundation for subsequent analysis and visualization.

### Graph Construction

To represent the metabolic interactions, we employed the NetworkX library in Python, a powerful tool for network analysis. The library provides a range of functionalities to construct and analyze graphs. We utilized the directed graph (DiGraph) class in NetworkX to capture the directional flow of metabolites and products in metabolic pathways.

### Node Creation

Using the table data obtained during the data collection phase, we created nodes for each component in the metabolic network. Nodes represented metabolites, proteins, enzymes, and other relevant entities. Each node was assigned a type, such as “metabolite,” “enzyme,” or “protein,” which allowed for easy categorization and subsequent visualization.

### Edge Creation

The edges in the graph represented the metabolic interactions between nodes. We defined three types of edges: metabolic_consumption, metabolic_reaction, and metabolic_production. These edge types captured different aspects of the metabolic processes. For example, metabolic_consumption edges denoted the consumption of a substrate, metabolic_reaction edges indicated biochemical reactions, and metabolic_production edges represented the production of a product. By assigning appropriate edge types, we could accurately depict the different interactions occurring within the metabolic network.

### Graph Visualization

After constructing the graph, we utilized the graph visualization capabilities of NetworkX and the matplotlib library to create clear and visually appealing representations of the metabolic interactions. We applied a layout algorithm, such as the spring layout, to position the nodes in a visually appealing manner. The colors of nodes and edges were customized based on their types to enhance the readability and interpretability of the visualizations.

According to Figure 6, we utilize “Biopython” library for accessing Entrez API. And the code can be formulated as follows.

**Figure 6:**
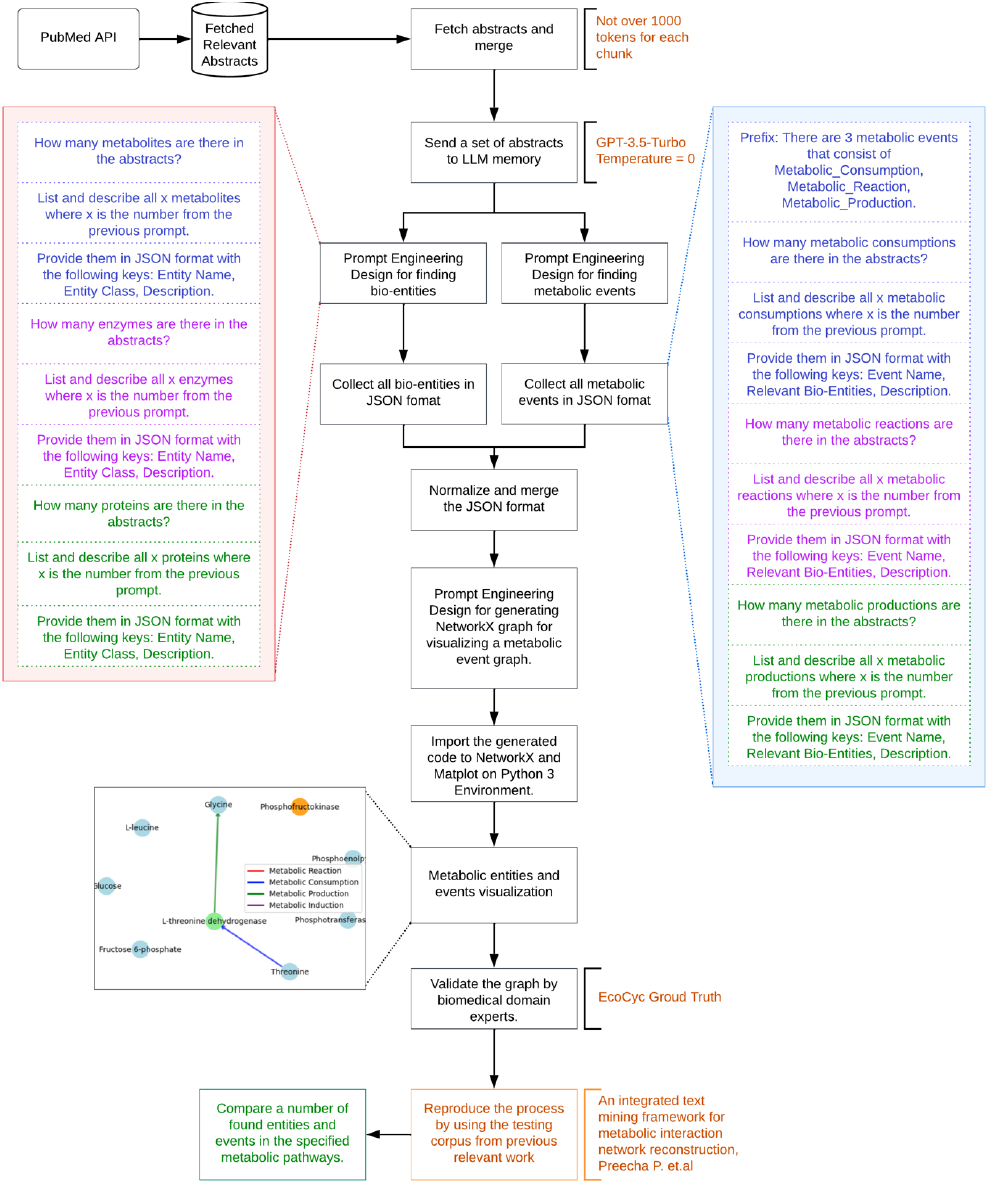
Metabolic Interaction Network Reconstruction Pipeline.

#python version 3.7x

pip install biopython

from Bio import Entrez

Before making any requests via Entrez API. It is required to provide our email address. This is done by registering on PubMed developer portal.

#python version 3.7x

Entrez.email = ““

By using the Entrez API to search for articles, we have to specify a search query and retrieve a list of matching article IDs using the esearch function from the Entrez module. For example, to search for articles related to a specific term can be coded as follows.

#python version 3.7x

search_term = “your_search_term”

handle = Entrez.esearch(db=“pubmed”, term=search_term) record = Entrez.read(handle)

article_ids = record[“IdList”]

Once we have the list of article IDs, we can retrieve the abstracts using the efetch function. Specify the database (pubmed), the list of article IDs, and the desired retmode (e.g., “text” for plain text) as follows.

#python version 3.7x

handle = Entrez.efetch(db=“pubmed”, id=article_ids, retmode=“text”, rettype=“abstract”)

abstracts = handle.read().split(‘\n\n’) for abstract in abstracts:

#Feed each abstract to GPT.

…

The retrieved abstracts will be stored in a list. So, we can process and utilize them for the next process that feed into GPT-3.5 Turbo model as in Figure 3.

**Table 1.**
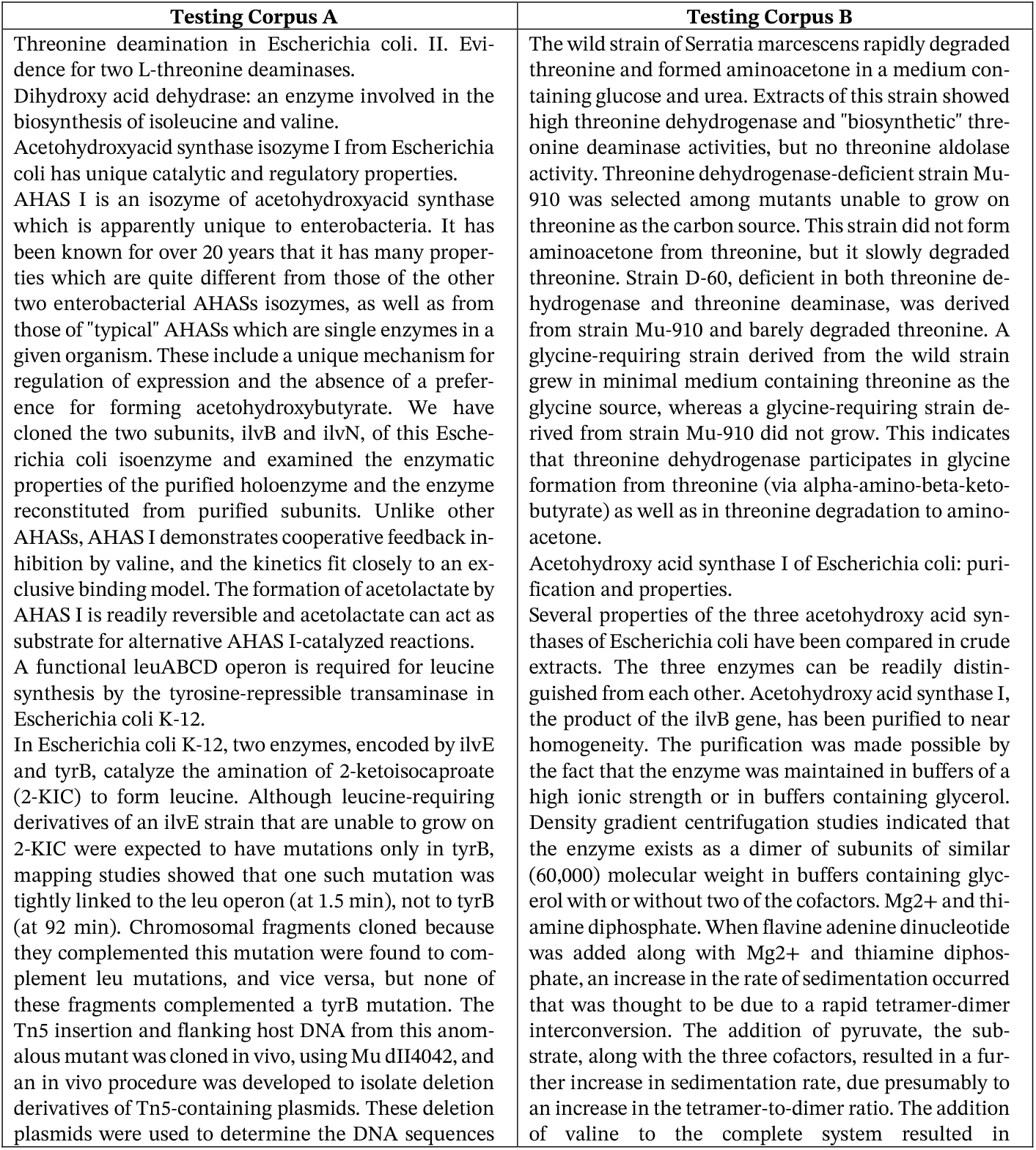
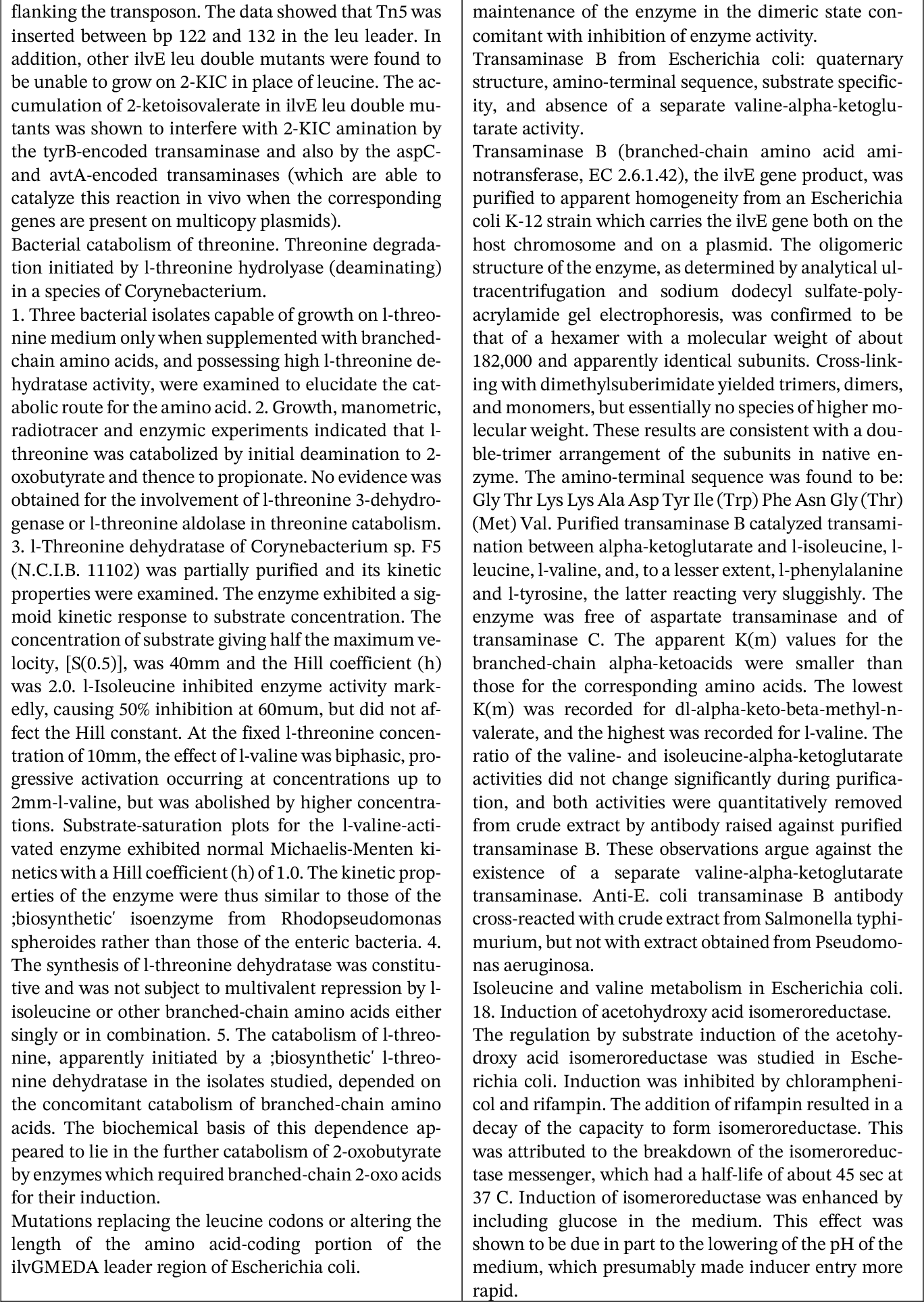
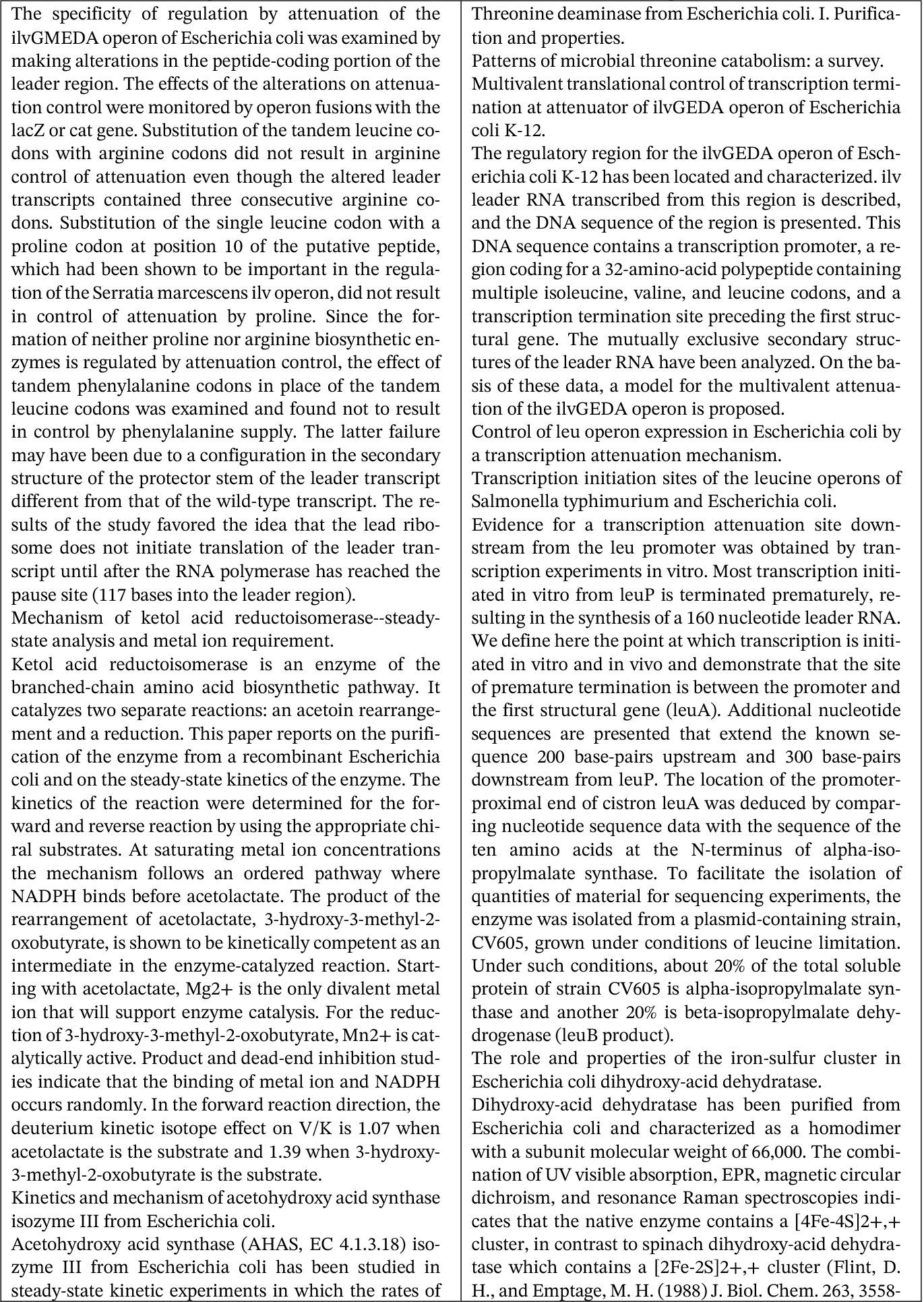
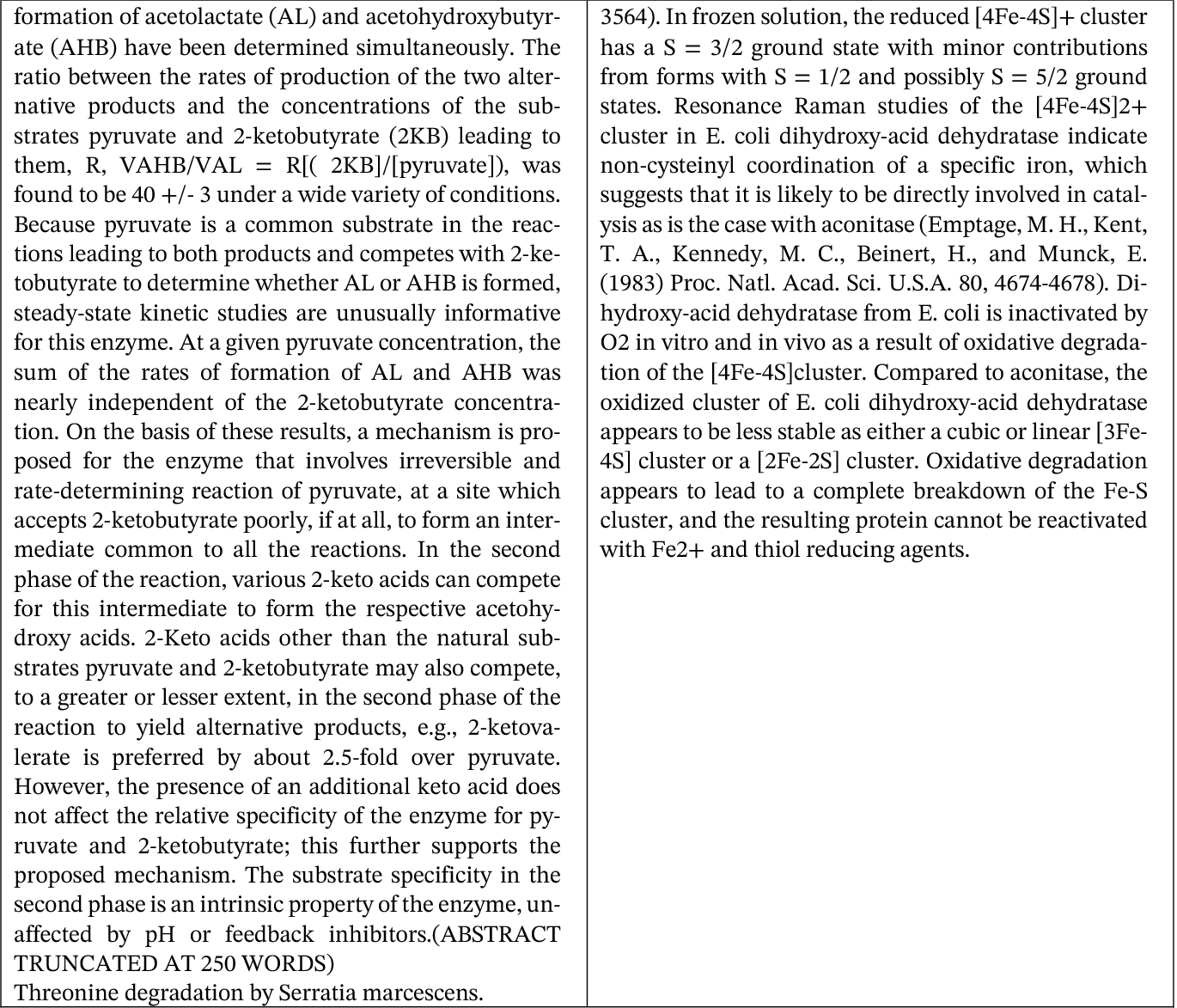
Testing abstracts from the previous relevant work [1].

| Testing Corpus A | Testing Corpus B |
|---|---|
| Threonine deamination in Escherichia coli. II. Evi-dence for two L-threonine deaminases.<br>Dihydroxy acid dehydrase: an enzyme involved in the biosynthesis of isoleucine and valine.<br>Acetohydroxyacid synthase isozyme I from Escherichia coli has unique catalytic and regulatory properties.<br>AHAS I is an isozyme of acetohydroxyacid synthase which is apparently unique to enterobacteria. It has been known for over 20 years that it has many proper-ties which are quite different from those of the other two enterobacterial AHASs isozymes, as well as from those of "typical" AHASs which are single enzymes in a given organism. These include a unique mechanism for regulation of expression and the absence of a prefer-ence for forming acetohydroxybutyrate. We have cloned the two subunits, ilvB and ilvN, of this Esche-richia coli isoenzyme and examined the enzymatic properties of the purified holoenzyme and the enzyme reconstituted from purified subunits. Unlike other AHASs, AHAS I demonstrates cooperative feedback in-hibition by valine, and the kinetics fit closely to an ex-clusive binding model. The formation of acetolactate by AHAS I is readily reversible and acetolactate can act as substrate for alternative AHAS I-catalyzed reactions.<br>A functional leuABCD operon is required for leucine synthesis by the tyrosine-repressible transaminase in Escherichia coli K-12.<br>In Escherichia coli K-12, two enzymes, encoded by ilvE and tyrB, catalyze the amination of 2-ketoisocaproate (2-KIC) to form leucine. Although leucine-requiring derivatives of an ilvE strain that are unable to grow on 2-KIC were expected to have mutations only in tyrB, mapping studies showed that one such mutation was tightly linked to the leu operon (at 1.5 min), not to tyrB (at 92 min). Chromosomal fragments cloned because they complemented this mutation were found to com-plement leu mutations, and vice versa, but none of these fragments complemented a tyrB mutation. The Tn5 insertion and flanking host DNA from this anom-alous mutant was cloned in vivo, using Mu dII4042, and an in vivo procedure was developed to isolate deletion derivatives of Tn5-containing plasmids. These deletion plasmids were used to determine the DNA sequences flanking the transposon. The data showed that Tn5 was inserted between bp 122 and 132 in the leu leader. In addition, other ilvE leu double mutants were found to be unable to grow on 2-KIC in place of leucine. The ac-cumulation of 2-ketoisovalerate in ilvE leu double mu-tants was shown to interfere with 2-KIC amination by the tyrB-encoded transaminase and also by the aspC- and avtA-encoded transaminases (which are able to catalyze this reaction in vivo when the corresponding genes are present on multicopy plasmids).<br>Bacterial catabolism of threonine. Threonine degrada-tion initiated by l-threonine hydrolyase (deaminating) in a species of Corynebacterium.<br>1. Three bacterial isolates capable of growth on l-threo-nine medium only when supplemented with branched-chain amino acids, and possessing high l-threonine de-hydratase activity, were examined to elucidate the cat-abolic route for the amino acid. 2. Growth, manometric, radiotracer and enzymic experiments indicated that l-threonine was catabolized by initial deamination to 2-oxobutyrate and thence to propionate. No evidence was obtained for the involvement of l-threonine 3-dehydro-genase or l-threonine aldolase in threonine catabolism. 3. l-Threonine dehydratase of Corynebacterium sp. F5 (N.C.I.B. 11102) was partially purified and its kinetic properties were examined. The enzyme exhibited a sig-moid kinetic response to substrate concentration. The concentration of substrate giving half the maximum ve-locity, [S(0.5)], was 40mm and the Hill coefficient (h) was 2.0. l-Isoleucine inhibited enzyme activity mark-edly, causing 50% inhibition at 60mum, but did not af-fect the Hill constant. At the fixed l-threonine concen-tration of 10mm, the effect of l-valine was biphasic, pro-gressive activation occurring at concentrations up to 2mm-l-valine, but was abolished by higher concentra-tions. Substrate-saturation plots for the l-valine-acti-vated enzyme exhibited normal Michaelis-Menten ki-netics with a Hill coefficient (h) of 1.0. The kinetic prop-erties of the enzyme were thus similar to those of the ;biosynthetic' isoenzyme from Rhodopseudomonas spheroides rather than those of the enteric bacteria. 4. The synthesis of l-threonine dehydratase was constitu-tive and was not subject to multivalent repression by l-isoleucine or other branched-chain amino acids either singly or in combination. 5. The catabolism of l-threo-nine, apparently initiated by a ;biosynthetic' l-threo-nine dehydratase in the isolates studied, depended on the concomitant catabolism of branched-chain amino acids. The biochemical basis of this dependence ap-peared to lie in the further catabolism of 2-oxobutyrate by enzymes which required branched-chain 2-oxo acids for their induction.<br>Mutations replacing the leucine codons or altering the length of the amino acid-coding portion of the ilvGMEDA leader region of Escherichia coli.<br>The specificity of regulation by attenuation of the ilvGMEDA operon of Escherichia coli was examined by making alterations in the peptide-coding portion of the leader region. The effects of the alterations on attenua-tion control were monitored by operon fusions with the lacZ or cat gene. Substitution of the tandem leucine co-dons with arginine codons did not result in arginine control of attenuation even though the altered leader transcripts contained three consecutive arginine co-dons. Substitution of the single leucine codon with a proline codon at position 10 of the putative peptide, which had been shown to be important in the regula-tion of the Serratia marcescens ilv operon, did not result in control of attenuation by proline. Since the for-mation of neither proline nor arginine biosynthetic en-zymes is regulated by attenuation control, the effect of tandem phenylalanine codons in place of the tandem leucine codons was examined and found not to result in control by phenylalanine supply. The latter failure may have been due to a configuration in the secondary structure of the protector stem of the leader transcript different from that of the wild-type transcript. The re-sults of the study favored the idea that the lead ribo-some does not initiate translation of the leader tran-script until after the RNA polymerase has reached the pause site (117 bases into the leader region).<br>Mechanism of ketol acid reductoisomerase--steady-state analysis and metal ion requirement.<br>Ketol acid reductoisomerase is an enzyme of the branched-chain amino acid biosynthetic pathway. It catalyzes two separate reactions: an acetoin rearrange-ment and a reduction. This paper reports on the purifi-cation of the enzyme from a recombinant Escherichia coli and on the steady-state kinetics of the enzyme. The kinetics of the reaction were determined for the for-ward and reverse reaction by using the appropriate chi-ral substrates. At saturating metal ion concentrations the mechanism follows an ordered pathway where NADPH binds before acetolactate. The product of the rearrangement of acetolactate, 3-hydroxy-3-methyl-2-oxobutyrate, is shown to be kinetically competent as an intermediate in the enzyme-catalyzed reaction. Start-ing with acetolactate, Mg2+ is the only divalent metal ion that will support enzyme catalysis. For the reduc-tion of 3-hydroxy-3-methyl-2-oxobutyrate, Mn2+ is cat-alytically active. Product and dead-end inhibition stud-ies indicate that the binding of metal ion and NADPH occurs randomly. In the forward reaction direction, the deuterium kinetic isotope effect on V/K is 1.07 when acetolactate is the substrate and 1.39 when 3-hydroxy-3-methyl-2-oxobutyrate is the substrate.<br>Kinetics and mechanism of acetohydroxy acid synthase isozyme III from Escherichia coli.<br>Acetohydroxy acid synthase (AHAS, EC 4.1.3.18) iso-zyme III from Escherichia coli has been studied in steady-state kinetic experiments in which the rates of formation of acetolactate (AL) and acetohydroxybutyr-ate (AHB) have been determined simultaneously. The ratio between the rates of production of the two alter-native products and the concentrations of the sub-strates pyruvate and 2-ketobutyrate (2KB) leading to them, R, VAHB/VAL = R[( 2KB]/[pyruvate]), was found to be 40 +/- 3 under a wide variety of conditions. Because pyruvate is a common substrate in the reac-tions leading to both products and competes with 2-ke-tobutyrate to determine whether AL or AHB is formed, steady-state kinetic studies are unusually informative for this enzyme. At a given pyruvate concentration, the sum of the rates of formation of AL and AHB was nearly independent of the 2-ketobutyrate concentra-tion. On the basis of these results, a mechanism is pro-posed for the enzyme that involves irreversible and rate-determining reaction of pyruvate, at a site which accepts 2-ketobutyrate poorly, if at all, to form an inter-mediate common to all the reactions. In the second phase of the reaction, various 2-keto acids can compete for this intermediate to form the respective acetohy-droxy acids. 2-Keto acids other than the natural sub-strates pyruvate and 2-ketobutyrate may also compete, to a greater or lesser extent, in the second phase of the reaction to yield alternative products, e.g., 2-ketova-lerate is preferred by about 2.5-fold over pyruvate. However, the presence of an additional keto acid does not affect the relative specificity of the enzyme for py-ruvate and 2-ketobutyrate; this further supports the proposed mechanism. The substrate specificity in the second phase is an intrinsic property of the enzyme, un-affected by pH or feedback inhibitors.(ABSTRACT TRUNCATED AT 250 WORDS)<br>Threonine degradation by Serratia marcescens. | The wild strain of Serratia marcescens rapidly degraded threonine and formed aminoacetone in a medium con-taining glucose and urea. Extracts of this strain showed high threonine dehydrogenase and "biosynthetic" thre-onine deaminase activities, but no threonine aldolase activity. Threonine dehydrogenase-deficient strain Mu-910 was selected among mutants unable to grow on threonine as the carbon source. This strain did not form aminoacetone from threonine, but it slowly degraded threonine. Strain D-60, deficient in both threonine de-hydrogenase and threonine deaminase, was derived from strain Mu-910 and barely degraded threonine. A glycine-requiring strain derived from the wild strain grew in minimal medium containing threonine as the glycine source, whereas a glycine-requiring strain de-rived from strain Mu-910 did not grow. This indicates that threonine dehydrogenase participates in glycine formation from threonine (via alpha-amino-beta-keto-butyrate) as well as in threonine degradation to amino-acetone.<br>Acetohydroxy acid synthase I of Escherichia coli: puri-fication and properties.<br>Several properties of the three acetohydroxy acid syn-thases of Escherichia coli have been compared in crude extracts. The three enzymes can be readily distin-guished from each other. Acetohydroxy acid synthase I, the product of the ilvB gene, has been purified to near homogeneity. The purification was made possible by the fact that the enzyme was maintained in buffers of a high ionic strength or in buffers containing glycerol. Density gradient centrifugation studies indicated that the enzyme exists as a dimer of subunits of similar (60,000) molecular weight in buffers containing glyc-erol with or without two of the cofactors. Mg2+ and thi-amine diphosphate. When flavine adenine dinucleotide was added along with Mg2+ and thiamine diphos-phate, an increase in the rate of sedimentation occurred that was thought to be due to a rapid tetramer-dimer interconversion. The addition of pyruvate, the sub-strate, along with the three cofactors, resulted in a fur-ther increase in sedimentation rate, due presumably to an increase in the tetramer-to-dimer ratio. The addition of valine to the complete system resulted in maintenance of the enzyme in the dimeric state con-comitant with inhibition of enzyme activity.<br>Transaminase B from Escherichia coli: quaternary structure, amino-terminal sequence, substrate specific-ity, and absence of a separate valine-alpha-ketoglu-tarate activity.<br>Transaminase B (branched-chain amino acid ami-notransferase, EC 2.6.1.42), the ilvE gene product, was purified to apparent homogeneity from an Escherichia coli K-12 strain which carries the ilvE gene both on the host chromosome and on a plasmid. The oligomeric structure of the enzyme, as determined by analytical ul-tracentrifugation and sodium dodecyl sulfate-poly-acrylamide gel electrophoresis, was confirmed to be that of a hexamer with a molecular weight of about 182,000 and apparently identical subunits. Cross-link-ing with dimethylsuberimidate yielded trimers, dimers, and monomers, but essentially no species of higher mo-lecular weight. These results are consistent with a dou-ble-trimer arrangement of the subunits in native en-zyme. The amino-terminal sequence was found to be: Gly Thr Lys Lys Ala Asp Tyr Ile (Trp) Phe Asn Gly (Thr) (Met) Val. Purified transaminase B catalyzed transami-nation between alpha-ketoglutarate and l-isoleucine, l-leucine, l-valine, and, to a lesser extent, l-phenylalanine and l-tyrosine, the latter reacting very sluggishly. The enzyme was free of aspartate transaminase and of transaminase C. The apparent K(m) values for the branched-chain alpha-ketoacids were smaller than those for the corresponding amino acids. The lowest K(m) was recorded for dl-alpha-keto-beta-methyl-n-valerate, and the highest was recorded for l-valine. The ratio of the valine- and isoleucine-alpha-ketoglutarate activities did not change significantly during purifica-tion, and both activities were quantitatively removed from crude extract by antibody raised against purified transaminase B. These observations argue against the existence of a separate valine-alpha-ketoglutarate transaminase. Anti-E. coli transaminase B antibody cross-reacted with crude extract from Salmonella typhi-murium, but not with extract obtained from Pseudomo-nas aeruginosa.<br>Isoleucine and valine metabolism in Escherichia coli. 18. Induction of acetohydroxy acid isomeroreductase.<br>The regulation by substrate induction of the acetohy-droxy acid isomeroreductase was studied in Esche-richia coli. Induction was inhibited by chlorampheni-col and rifampin. The addition of rifampin resulted in a decay of the capacity to form isomeroreductase. This was attributed to the breakdown of the isomeroreduc-tase messenger, which had a half-life of about 45 sec at 37 C. Induction of isomeroreductase was enhanced by including glucose in the medium. This effect was shown to be due in part to the lowering of the pH of the medium, which presumably made inducer entry more rapid.<br>Threonine deaminase from Escherichia coli. I. Purifica-tion and properties.<br>Patterns of microbial threonine catabolism: a survey.<br>Multivalent translational control of transcription termi-nation at attenuator of ilvGEDA operon of Escherichia coli K-12.<br>The regulatory region for the ilvGEDA operon of Esch-erichia coli K-12 has been located and characterized. ilv leader RNA transcribed from this region is described, and the DNA sequence of the region is presented. This DNA sequence contains a transcription promoter, a re-gion coding for a 32-amino-acid polypeptide containing multiple isoleucine, valine, and leucine codons, and a transcription termination site preceding the first struc-tural gene. The mutually exclusive secondary struc-tures of the leader RNA have been analyzed. On the ba-sis of these data, a model for the multivalent attenua-tion of the ilvGEDA operon is proposed.<br>Control of leu operon expression in Escherichia coli by a transcription attenuation mechanism.<br>Transcription initiation sites of the leucine operons of Salmonella typhimurium and Escherichia coli.<br>Evidence for a transcription attenuation site down-stream from the leu promoter was obtained by tran-scription experiments in vitro. Most transcription initi-ated in vitro from leuP is terminated prematurely, re-sulting in the synthesis of a 160 nucleotide leader RNA. We define here the point at which transcription is initi-ated in vitro and in vivo and demonstrate that the site of premature termination is between the promoter and the first structural gene (leuA). Additional nucleotide sequences are presented that extend the known se-quence 200 base-pairs upstream and 300 base-pairs downstream from leuP. The location of the promoter-proximal end of cistron leuA was deduced by compar-ing nucleotide sequence data with the sequence of the ten amino acids at the N-terminus of alpha-iso-propylmalate synthase. To facilitate the isolation of quantities of material for sequencing experiments, the enzyme was isolated from a plasmid-containing strain, CV605, grown under conditions of leucine limitation. Under such conditions, about 20% of the total soluble protein of strain CV605 is alpha-isopropylmalate syn-thase and another 20% is beta-isopropylmalate dehy-drogenase (leuB product).<br>The role and properties of the iron-sulfur cluster in Escherichia coli dihydroxy-acid dehydratase.<br>Dihydroxy-acid dehydratase has been purified from Escherichia coli and characterized as a homodimer with a subunit molecular weight of 66,000. The combi-nation of UV visible absorption, EPR, magnetic circular dichroism, and resonance Raman spectroscopies indi-cates that the native enzyme contains a [4Fe-4S]2+,+ cluster, in contrast to spinach dihydroxy-acid dehydra-tase which contains a [2Fe-2S]2+,+ cluster (Flint, D. H., and Emptage, M. H. (1988) J. Biol. Chem. 263, 3558-3564). In frozen solution, the reduced [4Fe-4S]+ cluster has a S = 3/2 ground state with minor contributions from forms with S = 1/2 and possibly S = 5/2 ground states. Resonance Raman studies of the [4Fe-4S]2+ cluster in E. coli dihydroxy-acid dehydratase indicate non-cysteinyl coordination of a specific iron, which suggests that it is likely to be directly involved in catal-ysis as is the case with aconitase (Emptage, M. H., Kent, T. A., Kennedy, M. C., Beinert, H., and Munck, E. (1983) Proc. Natl. Acad. Sci. U.S.A. 80, 4674-4678). Di-hydroxy-acid dehydratase from E. coli is inactivated by O2 in vitro and in vivo as a result of oxidative degrada-tion of the [4Fe-4S]cluster. Compared to aconitase, the oxidized cluster of E. coli dihydroxy-acid dehydratase appears to be less stable as either a cubic or linear [3Fe-4S] cluster or a [2Fe-2S] cluster. Oxidative degradation appears to lead to a complete breakdown of the Fe-S cluster, and the resulting protein cannot be reactivated with Fe2+ and thiol reducing agents. |

**Table 2.** Result validation by using GPT-3.5-Turbo Model.

|  |  |
|---|---|
| **Prompt :** | You are a biomedical data expert.<br>Ensure that you only analyze from my following abstracts delimited by triple backticks.<br>Don't make it rush, take your time to get the best solution.<br>There are 3 types of metabolic events including "Metabolic Consumption", "Metabolic Reaction", and "Metabolic Production".<br>Follow this instruction:<br>1. Check the result delimited by triple backticks that can be found in the abstracts.<br>2. If all entities are found in the abstracts, answer "Fully matched".<br>3. If some entities are not found in the abstracts, answer "Not matched" and show the list of not found entities. |
| **Response :** | Fully matched. |

## Results & Discussion

In this experiment, we use the testing corpus from previous work that includes 18 PubMed abstracts. First, we split the corpus into 2 corpora because GPT-3.5-Turbo model accepts only 4,000 tokens for analysis per time.

As in Figure 5, we have to optimize prompts iteratively, and his is the first prompt we use.

prompt = f” ” ”

You are a biomedical data expert. \

There are 2 main tasks for you and you need to make sure that a final result you \ provide is completely based on my information, not from other sources. \

Task 1: the task is to recognize biomedical entities from the provided text that consist of metabolites, enzymes, \ and proteins into a table with 2 headers that consist of “Entity Name” and “Entity Type”. \

Task 2: the task is to recognize metabolic event types from the provided text that consist of metabolic_reaction, \ metabolic_consumption, metabolic_production. And display its relevant entities in a table \

with 3 headers that consist of “Event Name”, “Event Type”, and “Relevant Entity” from Task 1 for each event.

“‘{abstracts}’”

As in Figure 7, we found that the LLM can response precisely for each task, but the result is difficult to be utilized for the further processes. So, we provide a specific format as HTML for obtaining a more proper result by using the prompt as follows.

**Figure 7:**
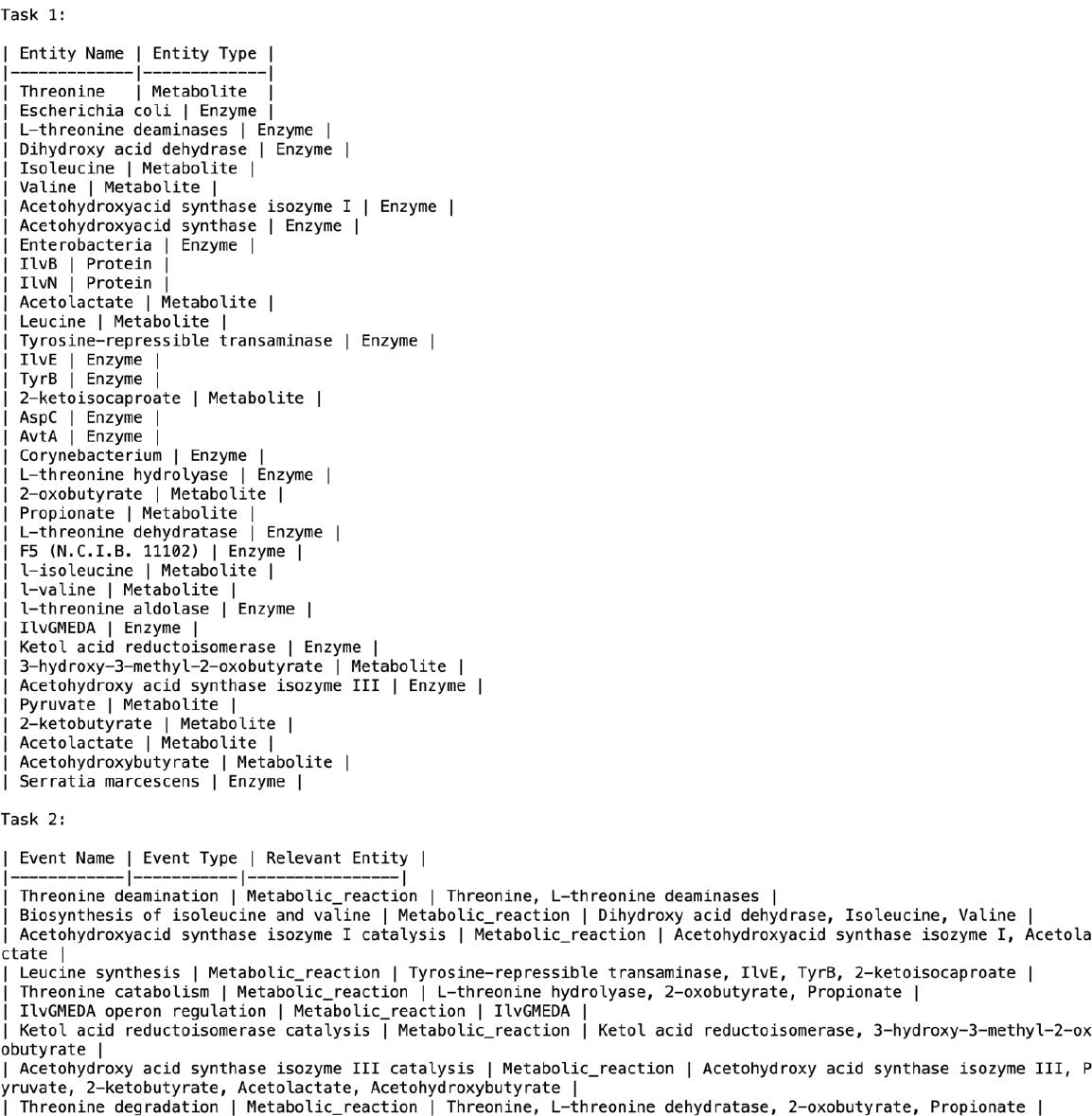
The result from the first prompt.

prompt = f”““

You are a biomedical data expert.

Ensure that you only analyze from my following abstracts delimited by triple backticks. Don’t make it rush, take your time to get the best result.

There are 3 bio-entity classes including “Metabolite”, “Enzyme”, and “Protein”. List all bio-entities from the abstracts based on the 3 classes.

Provide the result in HTML format with the following 3 headers: Entity Name, Entity Class, Description.

For the description, you need to describe about the entity name and the class.

Abstracts:

“‘{abstracts}’”

As a result, after optimizing the prompt, we obtain the result in HTML table format.

<h2>Metabolite</h2>

<table>

<tr>

<th>Entity Name</th>

<th>Entity Class</th>

<th>Description</th>

</tr>

<tr>

<td>l-threonine</td>

<td>Metabolite</td>

<td>An amino acid that is catabolized by initial deamination to 2-oxobutyrate and thence to propionate.</td>

</tr>

As in Figure 8, the result is separated into 3 tables by using Entity Class. Each table has 3 headers consisting of Entity Name, Entity Class, and Description. For Description, the model explains that in each row, why it chooses that word to be that entity class.

**Figure 8:**
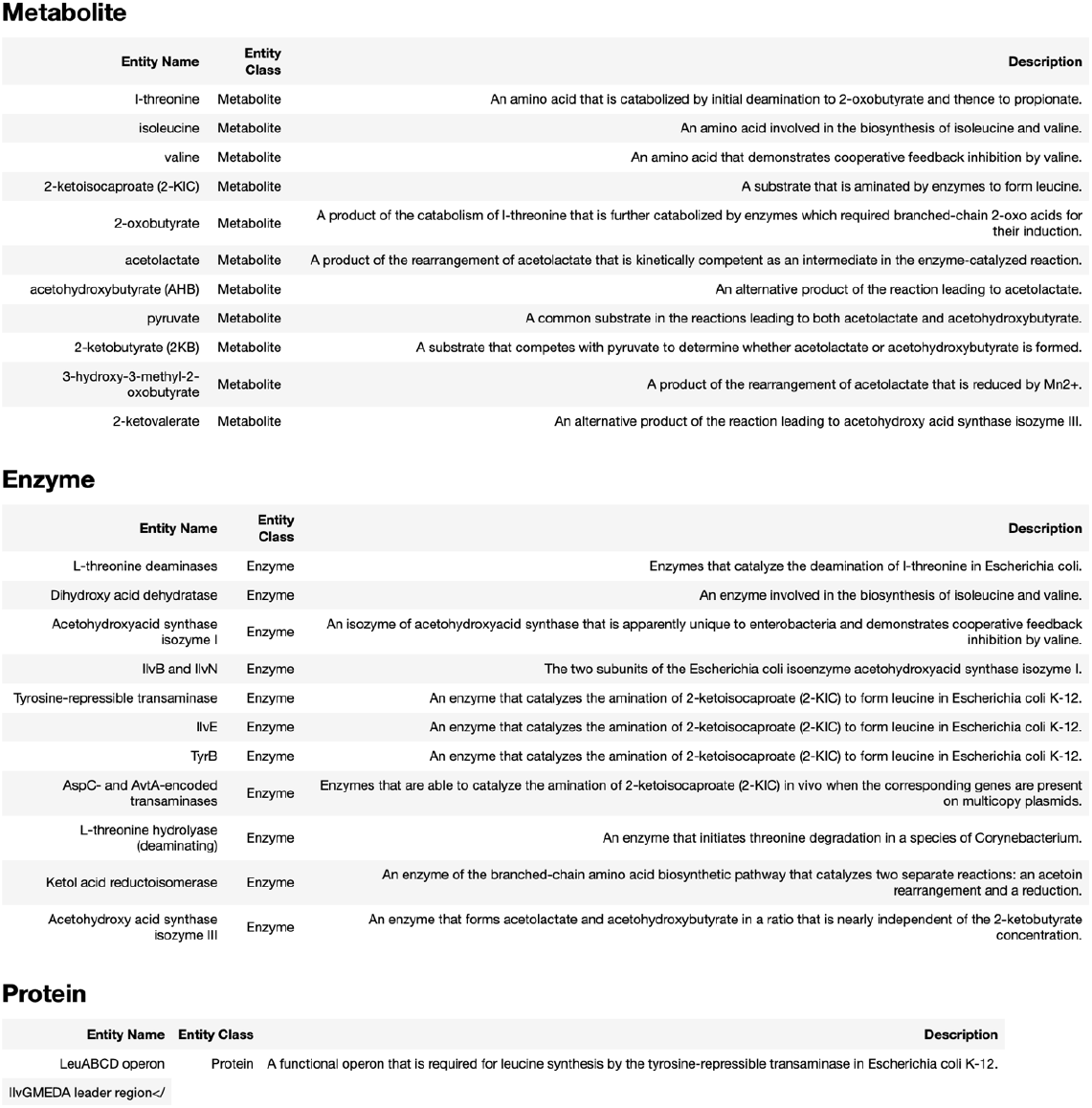
The result after optimizing the prompt.

**Figure 9:**
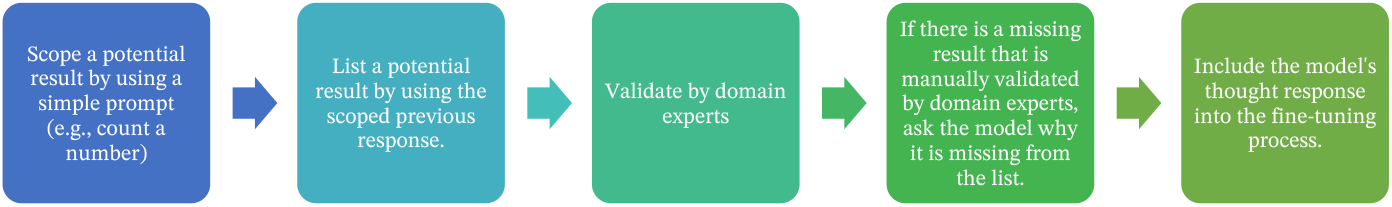
Fine-tuning process by using model response feedback.

However, although we configure the model parameter “Temperature” equals to “0” for preventing from hallucination, but the result seems not stable. For example, amounts of metabolites, enzymes as well as protein might not be the same in each execution time. Therefore, we have to fine-tune the prompt to be more specified for responding a more specified result.

As in Figure 6, we get started from letting the model counts a number of each entity class.

prompt = f’”

You are a biomedical data expert.

Ensure that you only analyze from my following abstracts delimited by triple backticks. Don’t make it rush, take your time to get the best result.

There are 3 bio-entity classes including “Metabolite”, “Enzyme”, and “Protein”. How many metabolites are there in the abstracts?

Abstracts:

“‘{abstracts}’”

As a result, the model responses as follows.

There are **<u>7 mentions</u>** of the bio-entity class “Metabolite” in the abstracts.

Thereafter, we optimize the previous prompt by using the result as follows.

prompt = f’”

You are a biomedical data expert.

Ensure that you only analyze from my following abstracts delimited by triple backticks. Don’t make it rush, take your time to get the best result.

There are 3 bio-entity classes including “Metabolite”, “Enzyme”, and “Protein”. List all <u>7 metabolites</u> from the abstracts.

Abstracts:

“‘{abstracts}’”

By using this prompt, the metabolite list is generated as below.

1. Isoleucine
2. Valine
3. Acetohydroxybutyrate
4. Acetolactate
5. 2-oxobutyrate
6. Propionate
7. 3-hydroxy-3-methyl-2-oxobutyrate

After manually validating by domain experts, they dive into the abstracts and found that “Pyruvate” is not included in the list. That means the model might not classify Pyruvate as a metabolite. Therefore, we create another prompt for enquiring about this issue as follows.

prompt = f’”

You are a biomedical data expert.

Ensure that you only analyze from my following abstracts delimited by triple backticks. Don’t make it rush, take your time to get the best result.

There are 3 bio-entity classes including “Metabolite”, “Enzyme”, and “Protein”. Is <u>Pyruvate</u> in the abstracts Metabolite? if not, please answer No and why.

Abstracts:

“‘{abstracts}’”

As in the abovementioned prompt, the model responses as below.

Yes, **<u>Pyruvate</u>** is mentioned in the abstracts, but it is **<u>not classified</u>** as a Metabolite. Pyruvate is a **<u>substrate</u>** or a r eactant in some of the enzymatic reactions described in the abstracts, but it is not explicitly classified as a Metabolite.

After taking a look by domain experts, we found that the model might not include some unstably classified entities in a final result. This problem is unable to be handled by Temperature parameter. Hence, we need to more clarify in the details of Pyruvate from GPT-3.5-Turbo model. To accomplish this task, we generate a prompt for the clarification as follow.

prompt = f’”

You are a biomedical data expert.

Ensure that you only analyze from my following abstracts delimited by triple backticks. Don’t make it rush, take your time to get the best result.

There are 3 bio-entity classes including “Metabolite”, “Enzyme”, and “Protein”. Why sometimes you classify Pyruvate as Metabolite, and some other time it is not?

Abstracts:

“‘{abstracts}’”

As the way that the LLM model thinks by using its very largely trained corpora, it can clarify reasonably as the following response which is also validated by domain experts.

<u>Pyruvate</u> is classified as a metabolite because it is a key intermediate in the metabolic pathway of glycolysis, which is the breakdown of glucose to produce energy. It is also involved in other metabolic pathways such as gluconeogenesis, the synthesis of amino acids, and the citric acid cycle. In some contexts, <u>pyruvate</u> may not be classified as a metabolite if it is being used as a substrate for an enzyme or if it is being produced as a byproduct of a specific reaction.

According to the response, we found that we have to fine-tune the current prompt by including more information about substrates as follows.

prompt = f’”

You are a biomedical data expert.

Ensure that you only analyze from my following abstracts delimited by triple backticks. Don’t make it rush, take your time to get the best result.

There are 3 bio-entity classes including “Metabolite”, “Enzyme”, and “Protein”.

How many metabolites <u>**including metabolic substrates**</u> are there in the abstracts?

Abstracts:

“‘{abstracts}’”

As a result, including metabolic substrates in the prompt can fulfill the unstable gap regarding the number of metabolites and the extracted metabolite list from the abstracts. And the response can be described as below.

There are a total of **<u>16 metabolites</u>** including **<u>metabolic substrates</u>** in the given abstracts. The metabolites are: threonine, isoleucine, valine, acetohydroxybutyrate, acetolactate, 2-ketoisocaproate, leucine, 2-ketoisovalerate, propionate, 2-oxobutyrate, phenylalanine, acetoin, 3-hydroxy-3-methyl-2-oxobutyrate, **<u>pyruvate</u>**, 2-ketobutyrate, and 2-ketovalerate.

To sum up regarding the result, the process of thought can be described as follows.

from tFor finding enzymes within the abstracts, we utilize the existing prompt that we use for finding metabolites as follows.

prompt = f’”

You are a biomedical data expert.

Ensure that you only analyze from my following abstracts delimited by triple backticks. Don’t make it rush, take your time to get the best result.

There are 3 bio-entity classes including “Metabolite”, “Enzyme”, and “Protein”. How many <u>**enzymes**</u> are there in the abstracts?

Abstracts:

“‘{abstracts}’”

According to the prompt, we still need to assign a number of bio-entity classes because we have to scope a boundary of the model’s thought.

As a result, the model provides feedback as following response.

There are a total of <u>**10 enzymes**</u> mentioned in the abstracts. To be more specific, the enzymes are: <u>**L-threonine deaminases, dihydroxy acid dehydrase, acetohydroxyacid synthase isozyme I, ilvB and ilvN, ilvE, tyrB, l-threonine hydrolyase, l-threonine dehydratase, ketol acid reductoisomerase, and acetohydroxy acid synthase isozyme III**</u>. Therefore, there are 10 enzymes in the abstracts.

After fine-tuning the prompt for finding enzymes with the specified number from the previous response, the fine-tuned prompt can be coded as follows.

prompt = f’”

You are a biomedical data expert.

Ensure that you only analyze from my following abstracts delimited by triple backticks. Don’t make it rush, take your time to get the best result.

There are 3 bio-entity classes including “Metabolite”, “Enzyme”, and “Protein”. List all <u>**10 enzymes**</u> from the abstracts.

Abstracts:

“‘{abstracts}’”

According to following response, it can be seen that the generated list is the same as in the previous response that describes all 10 enzyme names following its amount.

List of <u>**10 enzymes**</u>:

1. L-threonine deaminases
2. Dihydroxy acid dehydrase
3. Acetohydroxyacid synthase isozyme I
4. IlvB and IlvN subunits of AHAS I
5. TyrB-encoded transaminase
6. IlvE-encoded transaminase
7. L-threonine hydrolyase
8. L-threonine dehydratase
9. Ketol acid reductoisomerase
10. Acetohydroxy acid synthase isozyme III

For finding proteins, we can find by using a proper prompt for enquiring a number of proteins within the abstracts as follows.

prompt = f’”

You are a biomedical data expert.

Ensure that you only analyze from my following abstracts delimited by triple backticks. Don’t make it rush, take your time to get the best result.

There are 3 bio-entity classes including “Metabolite”, “Enzyme”, and “Protein”. How many <u>**proteins**</u> are there in the abstracts?

Abstracts:

“‘{abstracts}’”

And here below is the response.

There are 16 proteins mentioned in the abstracts.

The prompt can be changed for listing all 16 proteins as follows.

prompt = f’”

You are a biomedical data expert.

Ensure that you only analyze from my following abstracts delimited by triple backticks. Don’t make it rush, take your time to get the best result.

There are 3 bio-entity classes including “Metabolite”, “Enzyme”, and “Protein”. List all **<u>16 proteins</u>** from the abstracts.

Abstracts:

“‘{abstracts}’”

However, we found that some protein names in the following list are in the enzyme list.

List of 16 proteins:

1. L-threonine deaminases
2. Dihydroxy acid dehydrase
3. Acetohydroxyacid synthase isozyme I
4. IlvB
5. IlvN
6. IlvE
7. TyrB
8. AspC
9. AvtA
10. L-threonine hydrolyase
11. L-threonine dehydratase
12. IlvG
13. IlvM
14. IlvE
15. Ketol acid reductoisomerase
16. Acetohydroxy acid synthase isozyme III

The reason is an enzyme can be defined as a protein. A protein is a superset, and it also depends on how its function is within a chemical procedure. For more information, we provide a prompt for asking the model that how it classify as below.

prompt = f’”

You are a biomedical data expert.

Ensure that you only analyze from my following abstracts delimited by triple backticks. Don’t make it rush, take your time to get the best result.

There are 3 bio-entity classes including “Metabolite”, “Enzyme”, and “Protein”. Why <u>**enzyme “L-threonine deaminases” is also defined as a protein**</u>?

Abstracts:

“‘{abstracts}’”

After validating the below response by our domain expert, we found that the answer is true.

Enzyme “<u>**L-threonine deaminases**</u>” is also defined as a protein because it is a type of protein that catalyzes the deamination of L-threonine to produce 2-ketobutyrate. This is evident in the abstract “Threonine deamination in Escherichia coli. II. Evidence for two L-threonine deaminases.” which describes the presence of two L-threonine deaminases in Escherichia coli.

We can validate the result by checking within the provided abstracts to ensure that the model doesn’t generate by itself, accordingly to the generative model limitation in part of “hallucination”.

Next, we utilize the entities result that consists of metabolites, enzymes, and proteins to generate a directed graph for visualizing metabolic network reconstruction. To achieve this task, we have to fine-tune a prompt iteratively as in figure 10 because it needs to be scoped for generating a Python code properly.

**Figure 10:**
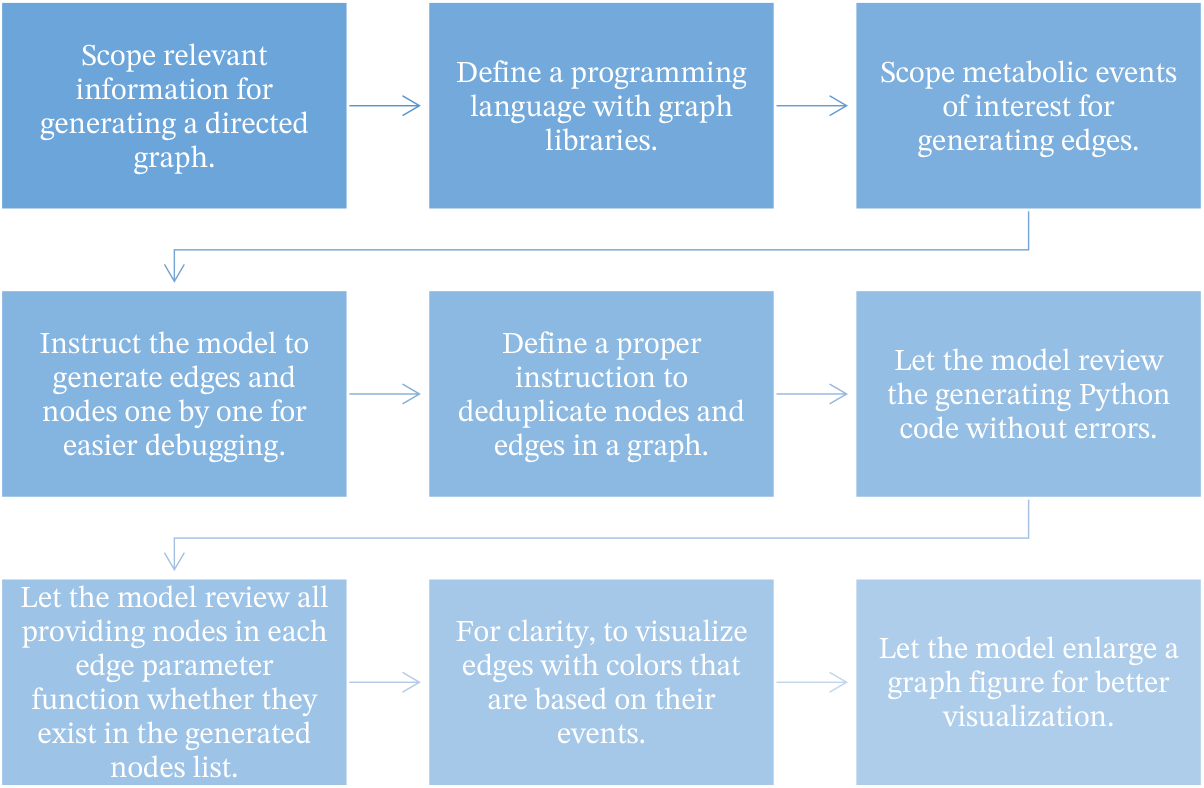
Fine-tuning iterative process for achieving a generative directed graph.

As in Figure 10, we can formulate a proper prompt as below.

You are a biomedical data expert.

Ensure that you only analyze from my following result delimited by triple backticks. Don’t make it rush, take your time to get the best solution.

There are 3 types of metabolic events including “Metabolic Consumption”, “Metabolic Reaction”, and “Metabolic Production”.

1. Based on the result and relevant information, provide a directed graph by using Python and NetworkX
2. Edges are metabolic events that consist of metabolic consumption, metabolic reaction, metabolic production. Create edges one by one, not in a tuple.
3. Ensure that there is no duplication within the graph.
4. Ensure that the python code completely generated without errors.
5. Ensure that nodes in the edge parameter are listed in the result.
6. Set edge colors based on events.
7. Enlarge the graph for viewing clearly and not too crowd.

According to the prompt, we ask for the generation of a directed graph using Python and NetworkX based on the provided result, which includes a list of metabolites, enzymes, and proteins. The graph should have edges representing metabolic events, such as metabolic consumption, metabolic reaction, and metabolic production.

The instructions emphasized avoiding duplication, ensuring all nodes are listed in the result, setting edge colors based on events, and enlarging the graph for clear visualization. To generate the Python code, the model first parses the provided result to extract the lists of metabolites, enzymes, and proteins.

Then, it initializes an empty directed graph using the NetworkX library. Next, the model adds the nodes to the graph with its respective node types (metabolite, enzyme, or protein). After that, it iterates through the metabolic events and added directed edges to the graph one by one, ensuring there were no duplicate edges. Finally, the graph is visualized using the matplotlib library, setting node colors, labels, and edge labels.

As in Figure 11, the graph is visualized by using the generated Python code. Some functions need to be reviewed by a software specialist because of a variety of library versions that may lead to a conflicted code. For the graph characteristics, it can be described as follows.

**Figure 11:**
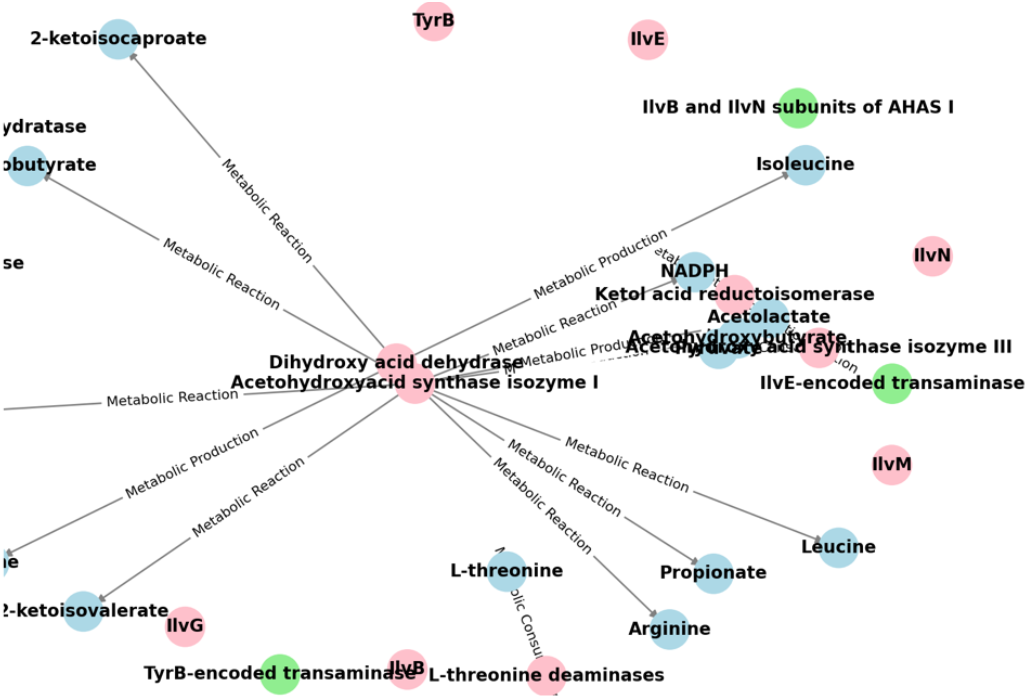
Metabolic network reconstruction from abstracts.

### Number of Metabolites

There are 16 metabolites in the graph. These metabolites include L-threonine, Isoleucine, Valine, Acetohydroxybutyrate, Acetolactate, 2-ketoisocaproate, Leucine, 2-ketoisovalerate, 2-oxobutyrate, Propionate, Arginine, Proline, NADPH, Acetolactate, Acetohydroxybutyrate, and Pyruvate.

### Number of Enzymes

There are 10 enzymes in the graph. These enzymes are L-threonine deaminases, Dihydroxy acid dehydrase, Acetohydroxyacid synthase isozyme I, IlvB and IlvN subunits of AHAS I, TyrB-encoded transaminase, IlvE-encoded transaminase, L-threonine hydrolyase, L-threonine dehydratase, and Ketol acid reductoisomerase.

### Number of Proteins

There are 16 proteins in the graph. These proteins include L-threonine deaminases, Dihydroxy acid dehydrase, Acetohydroxyacid synthase isozyme I, IlvB, IlvN, IlvE, TyrB, AspC, AvtA, L-threonine hydrolyase, L-threonine dehydratase, IlvG, IlvM, IlvE, Ketol acid reductoisomerase, and Acetohydroxy acid synthase isozyme III.

### Number of Metabolic Events

There are 25 metabolic events in the graph. These events consist of metabolic consumption, metabolic reaction, and metabolic production. Each event represents a directed edge connecting two nodes in the graph, indicating the relationship between a metabolite and an enzyme or protein.

In a directed graph, potential paths are sequences of nodes connected by directed edges, where each edge represents a specific metabolic event. The paths can start from any metabolite node and traverse through enzymes and proteins to reach other metabolite nodes. That means it depends on domain experts where they would like to explore. According to the graph visualization, before assigning metabolic events for the edges, the model must clarify every potential path.

### Metabolite to Enzyme Path

A path that starts from a metabolite node, connects to an enzyme node, and may continue to other enzymes or proteins. This path represents the involvement of an enzyme in metabolite conversion or production.

### Enzyme to Metabolite Path

A path that starts from an enzyme node, connects to other enzymes or proteins, and ends at a metabolite node. This path represents the utilization or production of a metabolite by enzymes.

### Metabolite to Protein Path

A path that starts from a metabolite node, connects to an enzyme node, and further connects to a protein node. This path represents the relationship between a metabolite and the protein associated with the enzyme involved in its metabolic conversion.

### Protein to Enzyme Path

A path that starts from a protein node, connects to an enzyme node, and may continue to other enzymes or proteins. This path represents the connection between a protein and the enzymes it interacts with in metabolic processes.

### Protein to Metabolite Path

A path that starts from a protein node, connects to an enzyme node, and ends at a metabolite node. This path represents the involvement of a protein in the metabolic conversion or production of a specific metabolite.

The key contribution of this graph is to visualize novel knowledge that is extracted from relevant published paper on PubMed. Domain experts can use this graph for comparing with the ground truth from EcoCyc for specific metabolic pathways. Potentially, it can be utilized for updating existing pathways more precisely and correctively.

## Conclusion

In this study, we used a large language model, GPT-3.5-Turbo, to analyze a testing corpus consisting of 18 PubMed abstracts. The corpus was split into two corpora, A and B, as the model accepts a maximum of 4,000 tokens for analysis per time. We aimed to extract information related to metabolites, enzymes, proteins, and their interactions to reconstruct a metabolic network.

After several iterations of prompt optimization, we obtained a comprehensive list of metabolites, enzymes, and proteins from the abstracts. We validated the results and found that all entities were fully matched with the provided abstracts. Using the extracted entities, we generated a directed graph for visualizing the metabolic network. The graph consisted of 16 metabolites, 10 enzymes, 16 proteins, and 25 metabolic events representing the relationships between these entities. The graph was visualized using Python and NetworkX, ensuring clear visualization by avoiding duplication and setting appropriate colors for nodes and edges.

The generated graph provides a visual representation of the metabolic network based on the information extracted from the PubMed abstracts. It can be used by domain experts for further analysis, comparison with existing metabolic pathways, and updating or refining those pathways. The graph serves as a valuable tool for exploring and understanding the metabolic interactions and pathways discussed in the abstracts. And here below list is our key contributions for this experiment.

### Automated Extraction of Metabolic Entities

The research demonstrates the capability of a language model, GPT-3.5-Turbo, to automatically extract metabolites, enzymes, and proteins from biomedical literature. This automated extraction can significantly speed up the process of identifying and cataloging important entities in metabolic research.

### Validation of Entity Extraction

The research provides a validation process to ensure the accuracy and completeness of the extracted entities. By comparing the extracted entities with the provided abstracts, researchers can assess the reliability of the language model’s results and identify any missing or mismatched entities.

### Metabolic Network Reconstruction

The research showcases the generation of a directed graph to visualize the metabolic network based on the extracted entities and their relationships. This network reconstruction provides a comprehensive overview of the metabolic interactions discussed in the PubMed abstracts and can serve as a foundation for further analysis and exploration.

### Visualization of Metabolic Pathways

The generated directed graph offers a visual representation of metabolic pathways, highlighting the connections between metabolites, enzymes, and proteins. This visualization aids in the understanding of complex metabolic processes and facilitates the identification of key components and their roles in metabolic pathways.

### Integration of Language Models and Network Analysis

The research demonstrates the integration of language models, such as GPT-3.5-Turbo, with network analysis tools like NetworkX. This combination allows for the automated extraction of information from text data and its transformation into a structured network representation, enabling efficient analysis and exploration of complex biological systems.

Overall, this research contributes to the field of biomedical informatics by showcasing the potential of large language models and network analysis techniques in extracting and visualizing metabolic information from scientific literature. It offers a valuable approach for researchers to gain insights into metabolic pathways and their components, ultimately advancing our understanding of biological systems.

## Acknowledgements

Thiptanawat Phongwattana thanks Royal Golden Jubilee (RGJ) for PhD. research fellowship. King Mongkut University of Technology Thonburi and IC2 lab are acknowledged for providing e-resources.

## Author Contributions

Thiptanawat P. and Jonathan H. Chan wrote the manuscript.

## Data Availability

No data was used for the research described in the article.

